# Comparative secretome of *Magnaporthe oryzae* identified proteins involved in virulence and cell wall integrity

**DOI:** 10.1101/2020.04.10.035139

**Authors:** Ning Liu, Linlu Qi, Manna Huang, Deng Chen, Changfa Yin, Yiying Zhang, Xingbin Wang, Guixin Yuan, Rui-Jin Wang, Jun Yang, You-Liang Peng, Xunli Lu

## Abstract

Plant fungal pathogens secrete numerous proteins into the apoplast at the plant–fungus contact sites to facilitate colonization. Only a few secreted proteins were functionally characterized in *Magnaporthe oryzae*, the fungal pathogen causing rice blast disease worldwide. ALG3 is an α-1, 3-mannosyltransferase function in N-glycan synthesis for secreted N-glycosylated proteins, and the Δ*alg3* mutants show strong defects in cell wall integrity and fungal virulence, indicating a potential effect on the secretion of multiple proteins. In this study, we compared the secretome of wild type and Δ*alg3* mutants, and identified 51 proteins that require ALG3 for proper secretion. These are predicted to be involved in metabolic processes, interspecies interactions, cell wall organization, and response to chemicals. The tested secreted proteins localized at the apoplast region surrounding the fungal infection hyphae. Moreover, the *N*-glycosylation of candidate proteins was significantly changed in the Δ*alg3* mutant, leading to the reduction of protein secretion and abnormal protein localization. Furthermore, we tested the function of two genes, one is a previously reported M. oryzae gene *Invertase 1* (*INV1*) encoding a secreted invertase, and the other one is a gene encoding an Acid mammalian chinitase (*AMCase*). The fungal virulence was significantly reduced and the cell wall integrity was altered in the Δ*inv1* and Δ*amcase* mutant strains. Elucidation of the comparative secretome of *M. oryzae* improves our understanding of the proteins that require ALG3 for secretion, and of their function in fungal virulence and cell wall integrity.

## Introduction

The fungal pathogen *Magnaporthe oryzae* causes rice blast disease, one of the most devastating diseases of cultivated rice (*Oryza sativa*) worldwide [1]. Due to its agronomic and scientific importance, *M. oryzae* has become a model fungus to study plant–pathogen interactions. As a hemibiotrophic fungus, *M. oryzae* undergoes an initial biotrophic infection stage and then switches to a necrotrophic stage [2, 3]. Once in contact with the plant surface, the conidium germinates and the germ tip forms a dome-shape appressorium on the leaf surface. The appressorium matures and develops a penetration peg to rupture the plant cuticle and invade the epidermal cell. After penetration, the fungus develops biotrophic invasive hyphae in the initial leaf cell and branches into neighboring cells. Eventually, the fungus generates a visible necrotic lesion with numerous, newly formed conidia ready for the next infection cycle [4]. *M. oryzae* mutants impaired in appressorium and/or infection hyphae formation also showed defects in pathogenicity, indicating that the formation of the fungal appressorium and biotrophic growth are essential for successful infection [5-10]. Therefore, exploring the molecular basis underlying these processes enable us to illustrate the underlined mechanisms of plant–fungus interactions and develop new anti-fungal strategies.

During the initial biotrophic infection stage, *M. oryzae* secretes numerous proteins into the plant–fungus contact sites to facilitate colonization. One class of secreted proteins are enzymes including cutinase2, endoglucanase, endo-β-1,4 xylanase and cellulases that break down the plant cell wall; the other class of secreted proteins are small effector proteins that suppress the host immune system or manipulate host metabolism [11-15]. *M. oryzae* effector proteins include cytoplasmic effectors and apoplastic effectors, which are secreted through distinct pathways [16]. The cytoplasmic effectors, such as PWL2 and BAS1, are secreted through exocyst components to the biotrophic interfacial complex, a specific plant membrane-rich structure associated with invasive hyphae. Many cytoplasmic effector proteins have been studied, especially the avirulent effector proteins including AvrPia, Avr1-CO39, AvrPita, AvrPik, and AvrPiz-t [17-23]. The apoplastic effectors, such as Slp1 and Bas4, are secreted from invasive hyphae into the extracellular compartment surrounding invasive hyphae via the conventional secretory pathway from the endoplasmic reticulum (ER) to the Golgi apparatus [8, 12, 16]. So far, only few apoplastic effector proteins have been characterized. The virulence effector Slp1 functions in host immune suppression by binding chitin oligosaccharides to avoid the chitin-induced plant immune response [12]. Since proteins secreted at early biotrophic fungal growth stages contribute to fungal pathogenicity, characterization of additional secreted proteins during invasive hyphae growth may help us to better understand the mechanisms of the rice–*M. oryzae* interaction.

*N*-glycosylation of protein is a post-translational modification commonly found in eukaryotic organisms. Many *N*-glycosylated proteins are plasma membrane-associated proteins or secreted proteins. The proper folding of these proteins relies on correct *N*-glycosylation process, which then related to a proper secretion route to their functional sites. *N*-glycosylation starts with the synthesis of the core oligosaccharide--NAcGlc_2_Man_9_Glc_3_--short for two N-acetylglucosamines (NAcGlc), nine mannoses (Man) and three glucose (Glc) molecules. After that, the core oligosaccharide is added to an asparagine residue (*N*) in the consensus sequence Asn-x-Ser/Thr (x is any amino acid apart from Proline). The *N*-glycosylation modification takes place in the ER and is modified in the Golgi apparatus, following the conventional secretion route for those target proteins. Defects in the early step of *N*-glycosylation results in an improper folding of target proteins, leading to protein breakdown through ER-associated degradation. Correct *N*-glycosylation is necessary for the proper function of secreted proteins in eukaryotes. For instance, impaired plant *N*-glycosylation or quality control for glycoprotein folding in the ER results in reduced plant resistance to bacterial pathogens. In pathogenic fungi, defects of *N*-glycosylation modification result in a reduction of fungal pathogenecity [24, 25]. In *M. oryzae*, loss of *ALG3*, an α-1, 3-mannosyltransferase functioning in core oligosaccharide synthesis, results in a significant reduction of fungal virulence and defects in fungal cell wall integrity. The apoplastic effector Slp1 was the target protein required ALG3 for its proper *N*-glycosylation [26]. Since Δ*alg3* exhibits a much stronger phenotype than that of the loss-of-function Δ*slp1* mutant, indicating apart from Slp1, other target proteins should also be affected in the Δ*alg3* mutant. Therefore, we attempted a proteomics approach to identify the other secreted proteins affected in the Δ*alg3* mutant, which should play essential roles in fungal virulence and cell wall integrity.

In this study, we used a comparative secretome analysis to identify secreted *N*-glycosylated proteins whose secretion required ALG3 function. Comparing the *in vitro* secretome of *M. oryzae* wild-type strain P131 and the knockout mutant Δ*alg3*, we found 51 proteins that were not secreted, or secreted in reduced amounts in the Δ*alg3* mutant. By testing 9 out of those 51 proteins, we confirmed that in the Δ*alg3* mutant, the *N*-glycosylation level of those proteins, as well as their localization in the infection hyphae were affected. We confirmed that two genes, *INV1* and *AMCase* function in fungal pathogenicity and cell wall integrity. Our study provided new insight about the role of secreted *N*-glycosylated proteins in fungal virulence and cell wall integrity.

## Results

### Identification of *M. oryzae* secreted proteins requiring ALG3 for proper secretion

In order to identify *N*-glycosylated proteins that require ALG3 for their secretion, we collected *M. oryzae* secreted proteins from the wild-type strain P131 and the knockout mutant Δ*alg3* grown in liquid growth medium (**Fig 1**A). *M. oryzae* may experience nutrient-deficient conditions when infecting susceptible plants; therefore, to better mimic the nutrient-deficient conditions during *M. oryzae* infection *in vitro*, we first compared the secreted proteins isolated from hyphae grown on nutrient-sufficient complete medium (CM) and nutrient-deficient minimal medium (MM). However, immunoblotting analysis using GFP fusion proteins found that Slp1-- a known effector protein whose secretion is regulated by ALG3-- only detected from *M. oryzae* grown in liquid CM but not from cultures grown in liquid MM (Supplemental Fig. 1A). Therefore, secreted proteins were collected from *M. oryzae* grown in liquid CM. Three replicates of extracted P131 and Δ*alg3* protein samples were separated by sodium dodecyl sulfate polyacrylamide gel electrophoresis (SDS-PAGE) and protein bands were clearly detected using silver staining (Supplemental Fig. 1B), indicating that those samples were suitable for subsequent analysis. The same amounts of secreted proteins were then used to the liquid chromatography with tandem mass spectrometry (LC-MS/MS) analysis for comparison.

**Figure 1.**
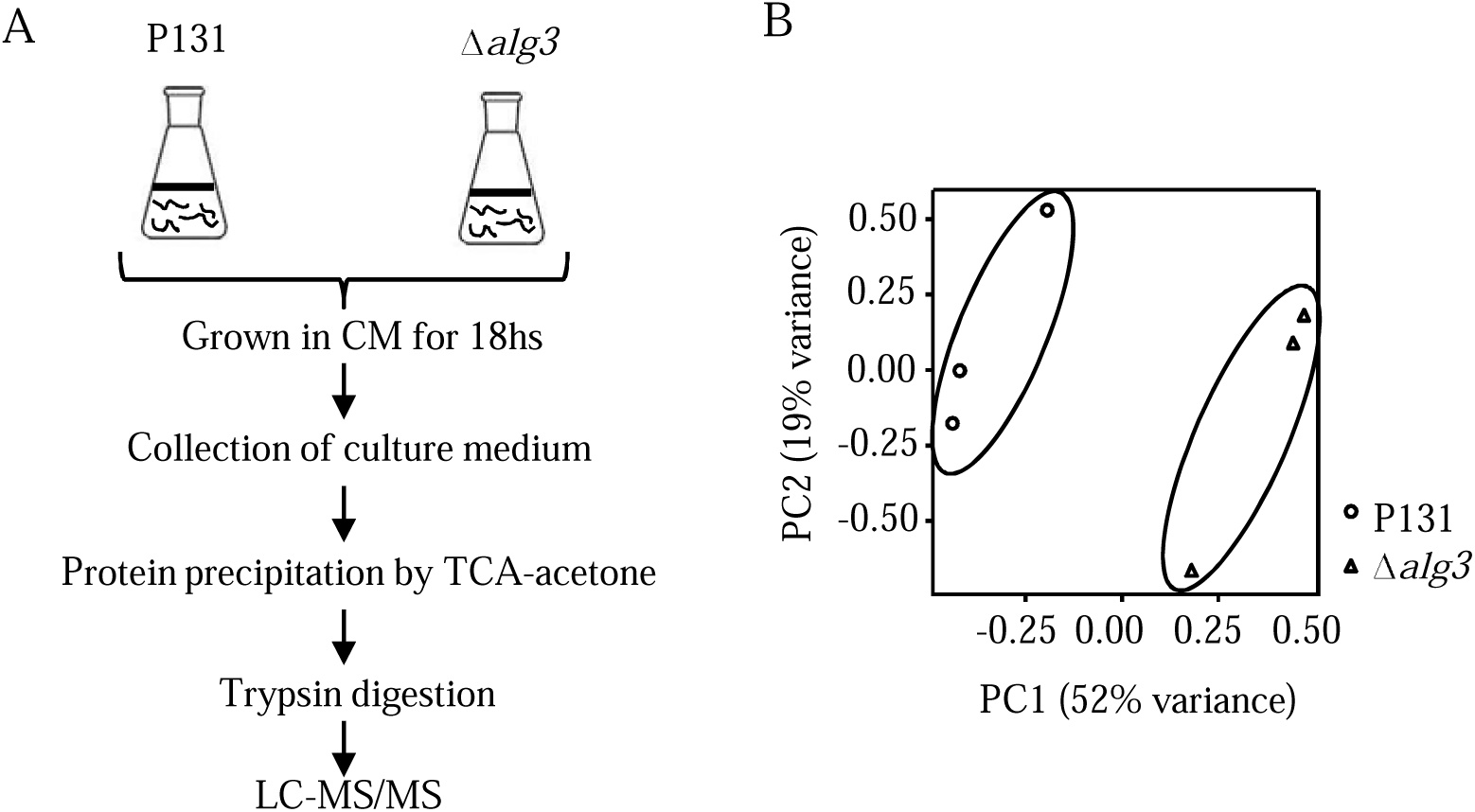
Quantitative secretome analysis of *Magnaporthe oryzae* secreted proteins from wild type P131 and Δ*alg3* mutant strains. **A**. Experimental strategy to identify secreted proteins of *M. oryzae* from liquid medium. **B**. PCA plot indicates the identified proteins from wild-type P131 strains and Δ*alg3* strains were grouped separately.

Using a label-free quantitative (LFQ) proteomic analysis, we identified 411 proteins from the wild-type P131 strain and 539 proteins from the Δ*alg3* mutant strain (Supplemental Table S1). The replication ratio was >74% for both samples (**Table 1** and Supplemental Table S1). We used Principal component analysis (PCA) to analyze the correlation and distribution of those secreted proteins from P131 and Δ*alg3* samples based on the protein intensity. In the PCA plot, P131 samples were clearly separated from Δ*alg3* samples (Fig. 1B), confirming that the isolated secreted proteins are different between P131 and Δ*alg3* strains.

**Table 1.**
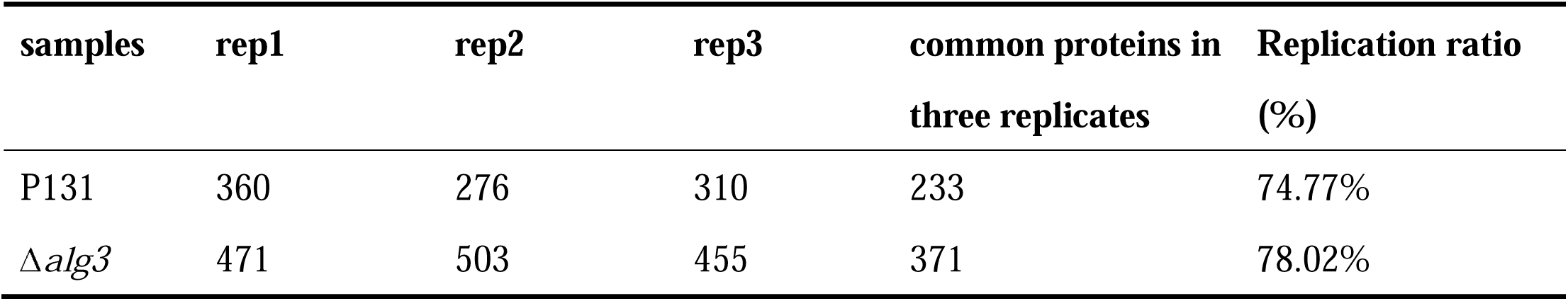
An overview of the identified secreted proteins of Mo from three biological replicates.

| **samples** | **rep1** | **rep2** | **rep3** | **common proteins in**<br>**three replicates** | **Replication ratio**<br>**(%)** |
| --- | --- | --- | --- | --- | --- |
| P131 | 360 | 276 | 310 | 233 | 74.77% |
| *Δalg3* | 471 | 503 | 455 | 371 | 78.02% |

Based on the protein LFQ intensities, we calculated the ratio of proteins in P131 to Δ*alg3* and selected proteins for which the fold-change ratio was >2.0 or <0.5 and the *p* value was <0.05. The identified proteins could be differentiated into three groups: 1) 51 proteins with a higher abundance in the secretome of P131 than in Δ*alg3*; 2) 295 proteins with an equal abundance in both secretomes; and 3) 210 proteins with a higher abundance in the secretome of Δ*alg3* than in P131 (**Table 2** and Supplemental Table S1). ALG3 is an *α*-1, 3-mannosyltransferase, which functions in the biosynthesis of the core oligosaccharide of N-glycan, and loss of ALG3 alters protein *N*-glycosylation modifications and reduces client protein steady-state levels, as shown previously for the effector protein Slp1. Therefore, proteins with significant lower abundance in Δ*alg3* (Group 1 proteins) are putative target proteins for *N*-glycosylation specifically require ALG3 for their secretion (**Table 3**).

**Table 2.** An overview of the number of proteins of different LFQ intensity groups.

| Groups | LFQ intensity | Total | SignalP | SecretomeP | TM |
| --- | --- | --- | --- | --- | --- |
| Group 1 | $P131 > \Delta alg3$ | 51 | 43 | 7 | 11 |
| Group 2 | $P131 = \Delta alg3$ | 295 | 128 | 69 | 22 |
| Group 3 | $P131 < \Delta alg3$ | 210 | 33 | 43 | 14 |

**Table 3.**
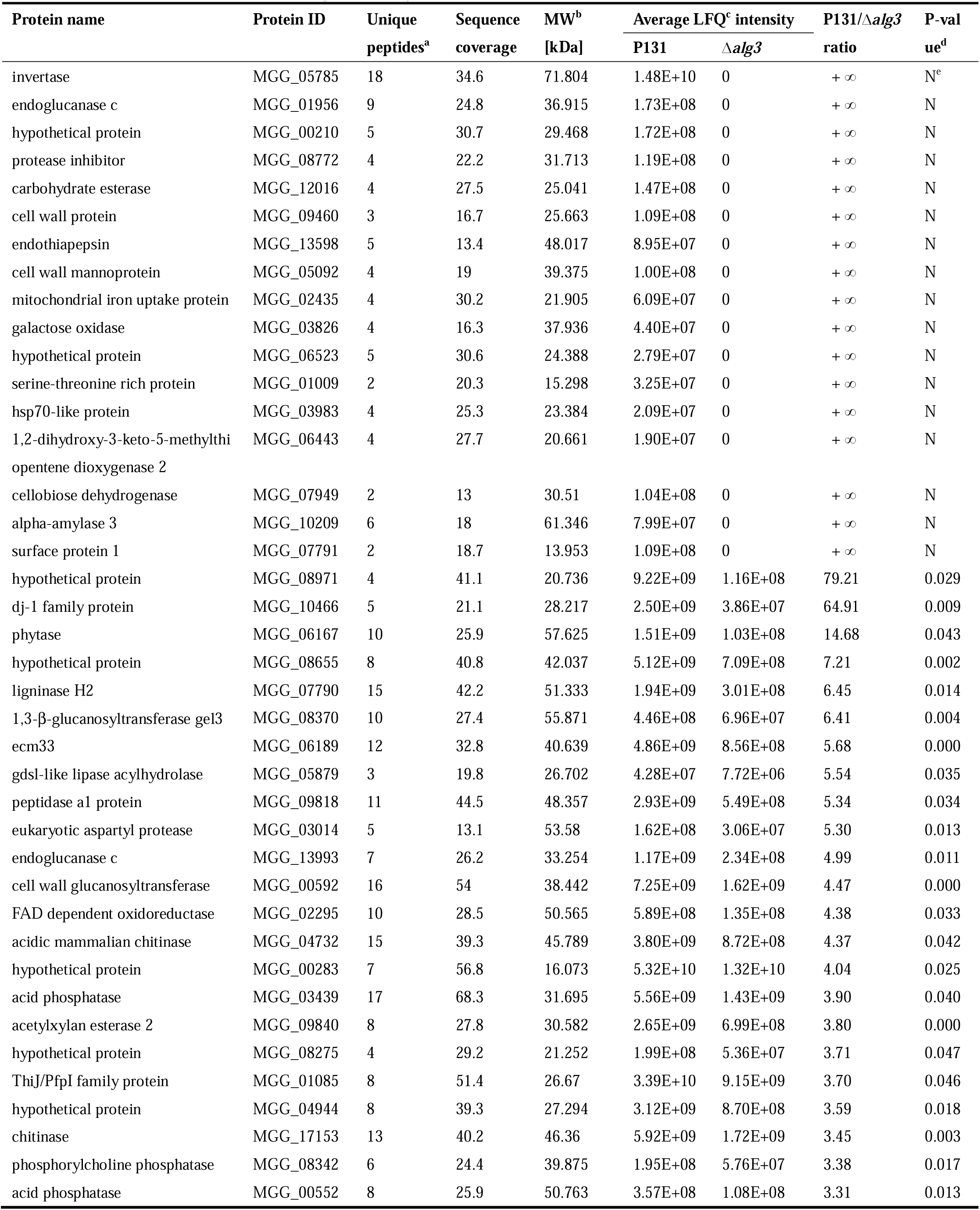
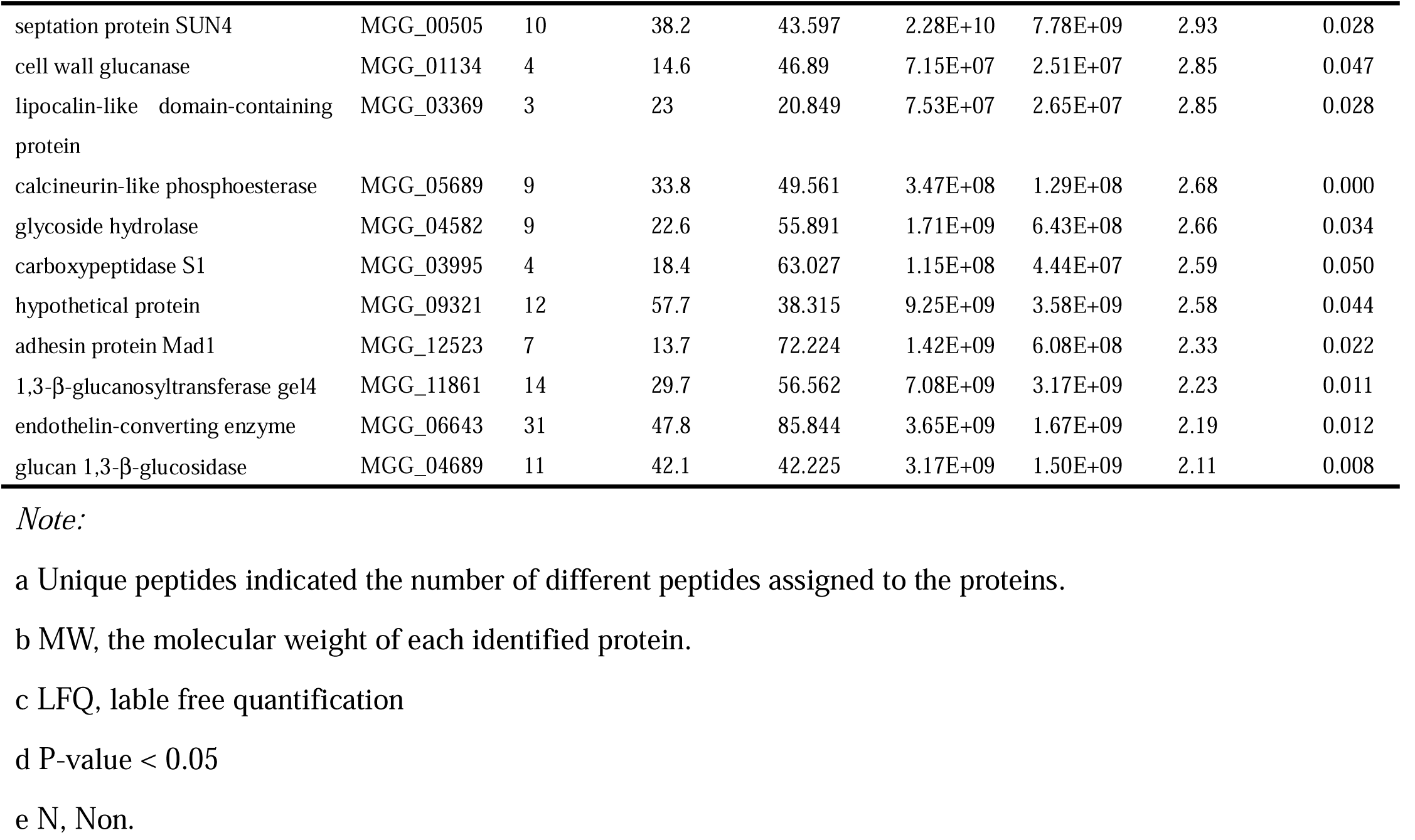
The secreted proteins regulated by ALG3.

We used SignalP (http://www.cbs.dtu.dk/services/SignalP-.4.1/), SecretomeP (http://www.cbs.dtu.dk/services/SecretomeP/), and Transmembrane Hidden Markov Model analyses (TMHMM, http://www.cbs.dtu.dk/services/TMHMM/) to validate the secreted nature of the proteins identified from secretome. SignalP predicts the N-terminal signal peptide for proteins secreted through the ER-Golgi pathway, SecretomeP predicts proteins non-classical secreted proteins, while TMHMM predicts possible transmembrane domains for membrane-anchored proteins. For Group 1 proteins, 43 proteins contain a SP and 7 proteins were predicted with SecretomeP, among them 11 proteins were predicted to have one TM domain. Therefore, 62.7% of Group 1 proteins are secreted proteins and 21.6% are membrane-localized proteins. Following the same calculation, 35.9% of Group 2 proteins are secreted proteins and 7.5% are membrane proteins; only 9.0% of Group 3 proteins are secretory proteins and 6.7% are membrane proteins. From the groups 3 proteins that specifically detected with higher abundance, a much smaller proportion are secreted proteins, indicating that many non-secreted proteins were abnormally secreted in the Δ*alg3* strain, possibly because the cell wall integrity was impaired in Δ*alg3* mutant; however, those proteins were not of interest in this study (Table 2 and Supplemental Table S2).

In further analysis, we focus on the 51 proteins listed in Table 3. We used *N*-GlycoSite analysis (http://www.cbs.dtu.dk/services/NetNGlyc/) to predict the possible *N*-linked glycosylation sites for them. Among them, 28 proteins (55%) have more than two predicted *N*-glycosylation sites, while 6 (12%), 11 (21%), and 6 (12%) proteins contain two, one, or no predicted *N*-glycosylation sites, respectively (**Figure 2**A and Supplemental Table 2). Thus, most proteins regulated by ALG3 contain predicted *N*-glycosylation sites.

**Figure 2.**
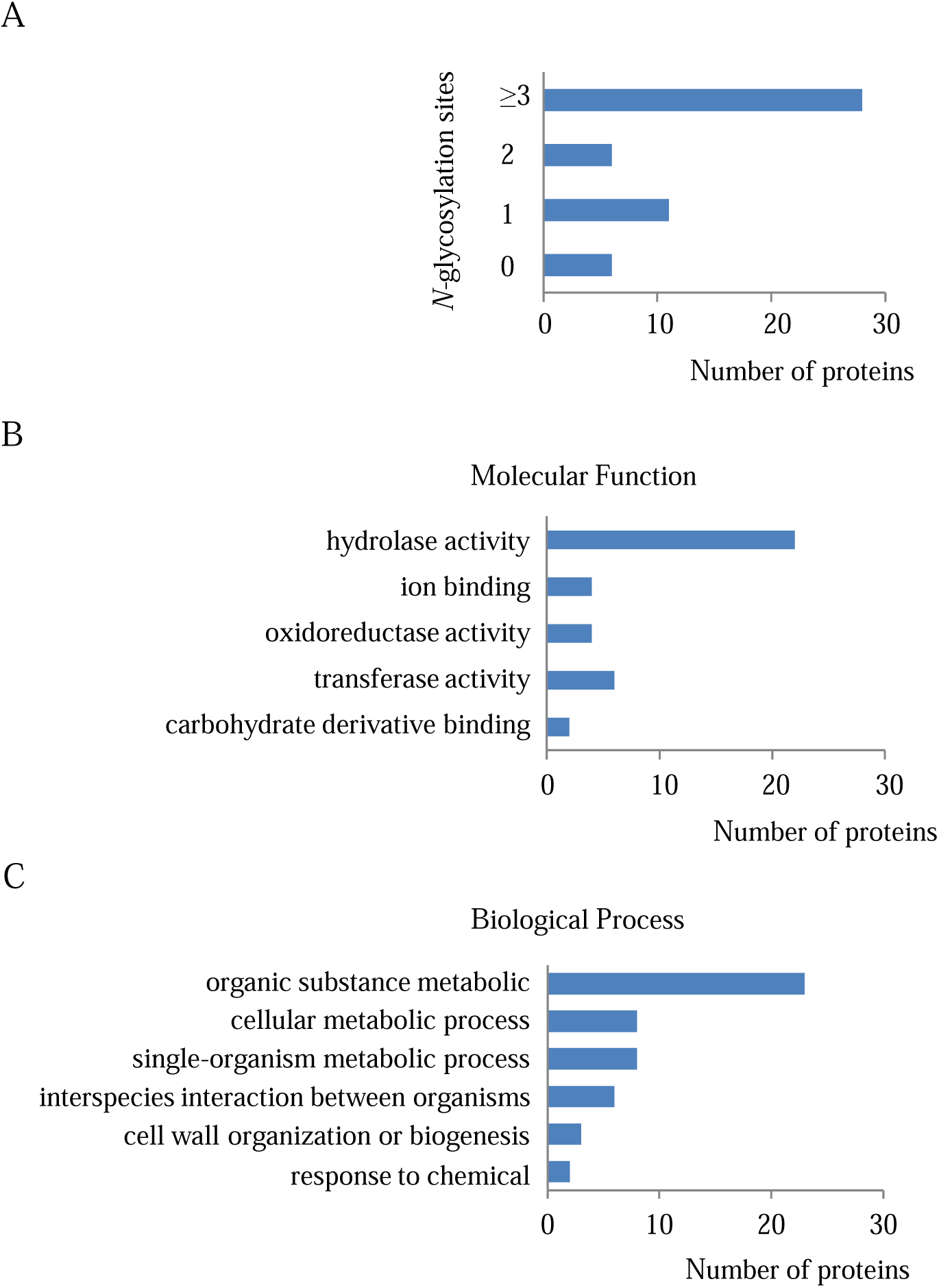
Bioinformatic analysis of secreted proteins requiring ALG3 for proper secretion. **A**. Predicted *N*-glycosylation sites using NetNGlyc alalysis for the secreted proteins requiring ALG3 for proper secretion. Gene Ontology analysis of those secreted proteins classified with molecular function category (**B)** and biological process category (**C)**.

Gene ontology enrichment analysis was performed for the 51 putative ALG3-regulated secreted proteins. Apart from eight hypothetical proteins with unknown functions, many proteins were detected with hydrolase, ion binding, and oxidoreductase activity (Figure 2B). Furthermore, many proteins were predicted to play a role in metabolic processes, eg: organic substance metabolites, cellular metabolites, and single-organism metabolic (Figure 2C), suggesting that those secreted proteins strongly influence *M. oryzae* metabolic processes. Other proteins were associated with cell wall organization, response to chemicals, or interaction between organisms (Figure 2C).

### Validation of secreted proteins regulated by ALG3

To further confirm the secretion of the identified proteins specifically regulated by ALG3, proteins were expressed in the *M. oryzae* P131 strain and Δ*alg3* strain, with a C-terminal GFP tag, under the control of the promotor from a strongly expressed *M. oryzae* gene (**Figure 3**A). We selected the following proteins for the validation tests: nine proteins including MGG_05785, MGG_01956, MGG_08772, MGG_09460, MGG_03826, MGG_10209, MGG_10466, MGG_00592 and MGG_04732 were identified with a significantly higher abundance in the medium of P131 compared with that of Δ*alg3* (Group 1 proteins, table 3), and one protein MGG_13764 identified at a similar level in the medium of both strains was chosen as a control sample for the secretion assay (Group 2 protein, Supplemental Table 2).

**Figure 3.**
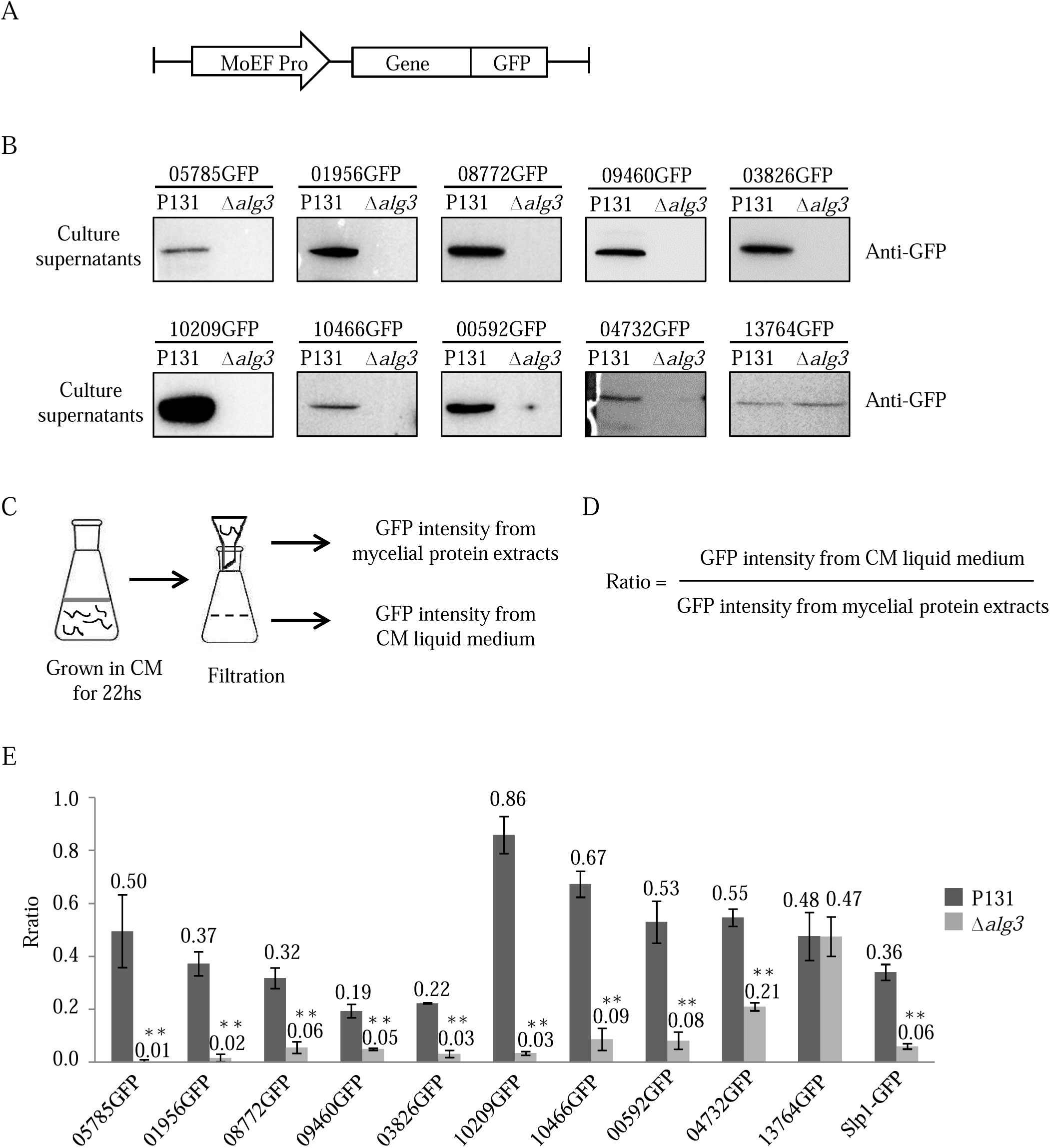
Secretion of the tested proteins requiring ALG3 for proper secretion. **A**. Schematic diagram of constructed vectors for tested genes. MoEF pro, *M oryzae* elongation factor promoter; Gene, genomic DNA sequences of target genes. Each constructed vector was transformed into wild-type P131 and Δ*alg3* strains. **B**. Immunoblotting analysis of tested proteins from culture supernatants from wild-type P131 and Δ*alg3* strains. Three replicates were performed for each strain, and one representative figure was shown. **C**. Experimental strategy to measure the GFP intensity from mycelial protein extracts and CM liquid culture. **D**. The GFP intensity ratio was calculated as the GFP intensity from 100μl CM medium versus that from 0.1g mycelium protein extracts. **E**. The GFP intensity ratio calculated for the tested secreted proteins in wild-type P131 and Δ*alg3* strains. Slp1, a known effector protein regulated by ALG3, was used as a positive control. Error bars denote standard deviations from three biological replicates. **, statistically significant differences between the wild type and mutants (P<0.01, Student’s t test).

We cultured various fungal strains in liquid CM and detect the level of proteins in the secreted CM medium using anti-GFP antibody. We found that 05785-GFP, 01956-GFP, 08772-GFP, 09460-GFP, 03826-GFP, 10209-GFP and 10466-GFP were detected in the CM medium of P131 strain but not of Δ*alg3* strain; while 00592-GFP and 04732-GFP were detected in the CM medium of P131 strain, and barely detectable in that of Δ*alg3* strain, and 13764-GFP were detected in the CM of both P131 and Δ*alg3* strains (Figure 3B). All the proteins were detectable in the mycelia protein extracts (see below), indicating efficient protein expression in those fungal strains. The immunoblotting assays confirmed our secretome data that tested secreted proteins from Group 1 require ALG3 for proper secretion.

We developed another method to directly test protein secretion in the liquid medium without a protein extraction step. The fluorescence signal from a functional GFP fusion can be easily measured using a microplate reader, and the fluorescence intensity correlates to the protein expression level (Figure 3C). Therefore, the liquid CM was loaded in the microplate reader and the fluorescence intensity was measured to detect secreted GFP fusion proteins. Considering that the same vector might integrate into different genomic regions in different strains, we could not directly compare the measured fluorescence intensity between P131 and Δ*alg3* strains harboring the same GFP fusion protein. Therefore, the fluorescence intensity of protein extracts from a certain weight (0.1g) of *M. oryzae* mycelia were also measured for normalization. For each strain, the fluorescence signals of the liquid CM were divided by that of mycelium protein extracts and calculated as a ratio to determine the level of secretion of each protein (Fig. 3D). To test whether this measurement works for secreted proteins, the fluorescence intensity of the P131 strain expressing control GFP alone was measured and the rate of secretion was calculated as 0.08, indicating that unfused GFP was not secreted into the liquid CM. Moreover, Slp1-GFP was secreted in the P131 strain at a rate of 0.36, while the nuclei-localized protein MoGRP1-GFP was secreted at a rate of 0.06, supporting the observation that Slp1-GFP but not MoGRP1-GFP was secreted into the liquid CM in the P131 strain [27] (Supplemental Fig. 2B). Furthermore, in the Δ*alg3* strain, Slp1-GFP was secreted at a rate of 0.06, confirming the finding that the secretion of Slp1-GFP is impaired in the Δ*alg3* strain. These controls confirm that this measurement and calculation is suitable to detect secreted GFP fusion proteins in *M. oryzae* strains.

Using the method described above, we validated that proteins encoded by MGG_05785, MGG_01956, MGG_08772, MGG_09460, MGG_03826, MGG_10209, MGG_10466, MGG_00592 were secreted in the liquid CM from the P131 strains, but were not secreted from the Δ*alg3* strains (Fig. 3E), indicating that those proteins were indeed secreted proteins regulated by ALG3. Protein MGG_04732 were secreted in the liquid CM of the P131 strains but with a reduced level in the liquid of Δ*alg3* strains. The control protein encoded by MGG_13764, were secreted in the liquid CM of both P131 and Δ*alg3* strains (Fig. 3E), further confirmed our secretome data on Group1 proteins.

We next tested whether those secreted proteins contain the *N*-glycosylation modification. Our previous study demonstrated that the client protein Slp1 were N-glycosylated with Man_5_GlcNAc_2_ (but not Man_9_GlcNAc_2_) in the Δ*alg3* mutants, and the size difference was observed using immunoblot analysis. We found that all the tested proteins differed in size between the P131 and Δ*alg3* strains (**Fig. 4**), indicating that complete *N*-glycosylation of the above-mentioned proteins occurs in the P131 strain, while only partial *N*-glycosylation takes place in the Δ*alg3* strain. We also applied the peptide *N*-glycosidase F (PNGase F) [28]--an amidase that cuts the GlcNAc group and the Asn residues from *N*-glycosylated proteins--and detected the de-glycosylated form of each protein in the P131 strain (Fig. 4). The different protein bands between the minus and plus PNGase F treatment in the Δ*alg3* strain suggested that the partial *N*-glycosylation of glycoproteins was also cleaved by PNGase F. Our findings revealed that the identified secretory proteins contained the *N*-glycosylation modification and depend on ALG3 for a complete *N*-glycosylation.

**Figure 4.**
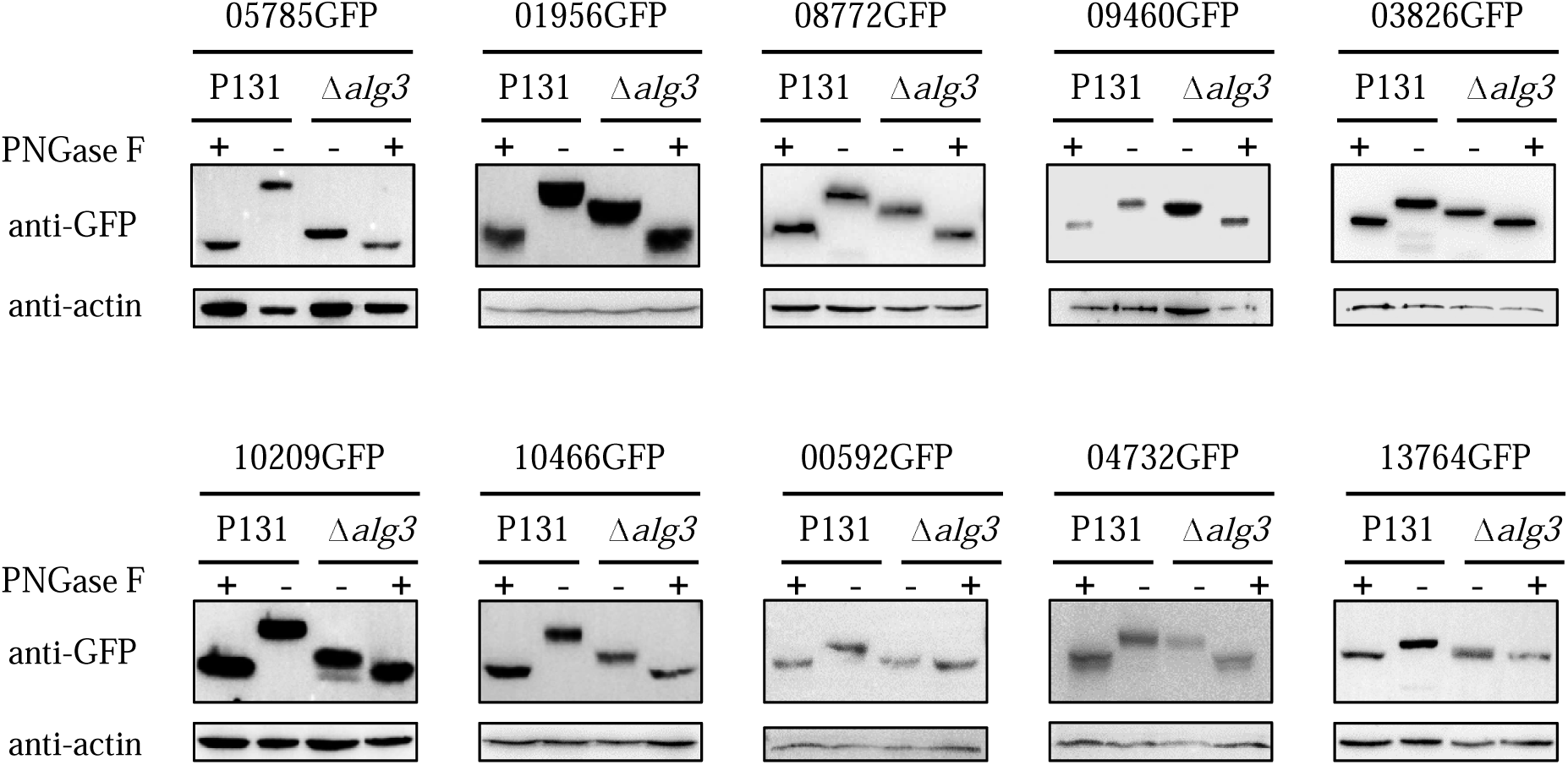
Assays for *N*-glycosylation of secreted proteins in the wild-type P131 and Δ*alg3* strains. Mycelial protein lysates with (+) or without (−) PNGase F treatment were immunoblotted with anti-GFP antibody. Protein loading control was detected using anti-Actin antibody.

Since those secreted proteins were identified from liquid medium during fungal vegetative growth, we wondered whether they also localized in the infection hypha during the plant–fungus interaction. We investigated their subcellular localization in the infection hyphae using a fluorescence microscope. In P131 strains transformed with GFP-labeled MGG_05785, MGG_01956, MGG_08772, MGG_09460, MGG_03826, MGG_10209, MGG_10466, MGG_00592, MGG_04732 and MGG_13764, the GFP signals were distributed in the cytoplasm and at the plant–fungus interface (**Fig. 5**), similar to Slp1-GFP in the P131 strain. In Δ*alg3* strains transformed with GFP-labeled MGG_05785, MGG_01956, MGG_08772, MGG_09460, MGG_03826, MGG_10209, MGG_10466, MGG_00592 and MGG_04732, the GFP fusion proteins were restricted in small dot-like structures within the infection hyphae and only a small portion of GFP signals located at the plant–fungus interface (Fig. 5), indicating that *N*-glycosylation is important for the subcellular localization of those proteins. In Δ*alg3* strains expressing MGG_13764-GFP, the fluorescent signals were similar to that in P131 strains, indicating that *N*-glycosylation modification is not essential for the subcellular localization of MGG_13764 protein.

**Figure 5.**
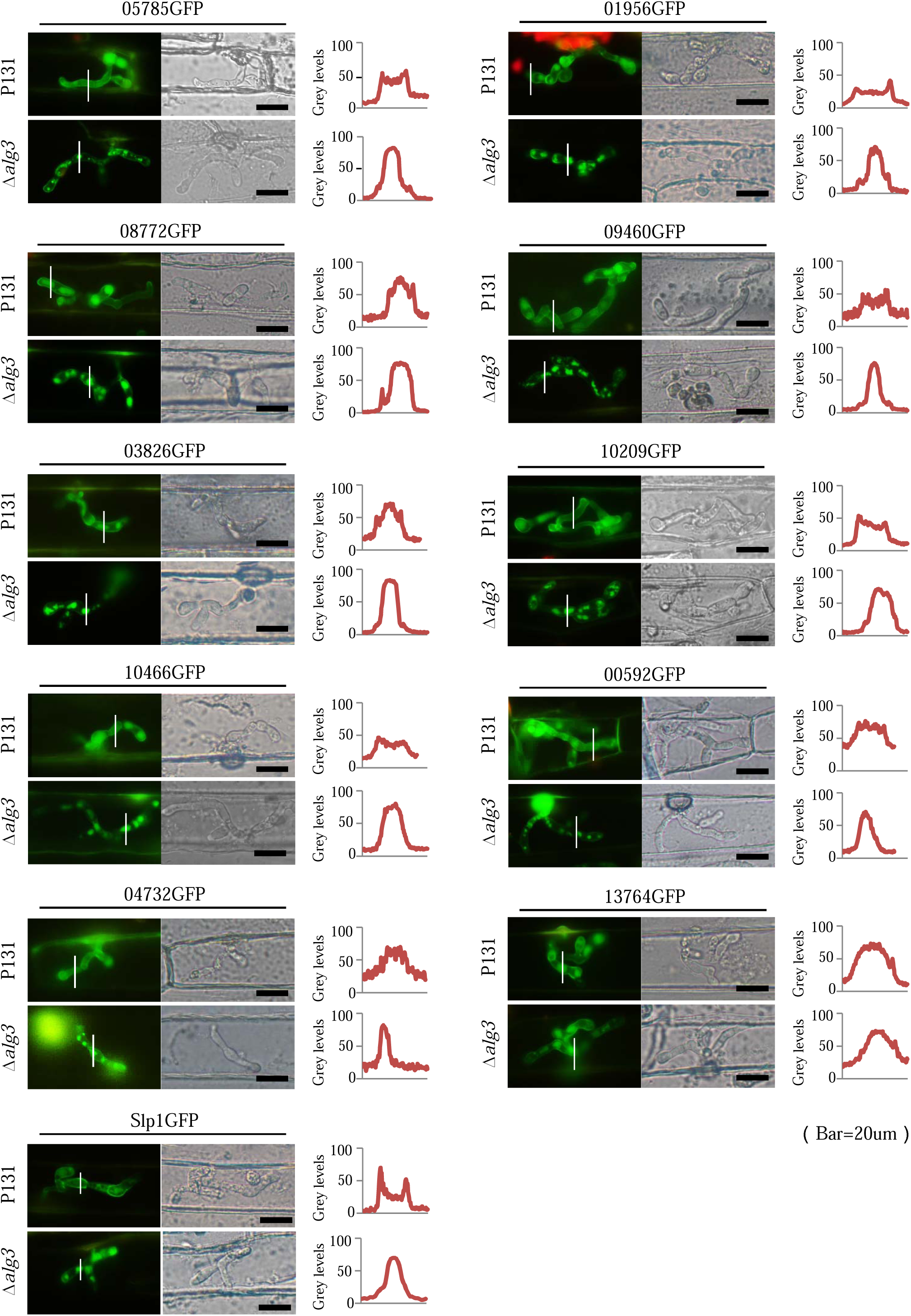
The localization of secreted proteins in the infected hyphae differs in wild-type P131 and Δ*alg3* strains. Left panel, Fluorescence signal is imaged in infected hyphae growing in barley epidermal cells for the tested proteins. White lines in the figures show the path for fluorescence intensity measurement. Middle panel, the brightfield image for the infection hyphae. Right panel, the fluorescence intensity measured using ImageJ. Scale bar=20 μm

### MGG_05785 is important for fungal pathogenicity and cell wall integrity

From the LC-MS data, INV1, encoded by *MGG_05785*, is the most abundant protein identified only in liquid CM of the P131 strain (Table 3). The *N*-glycosylation modification of INV1 was confirmed in the mycelium proteins of *M. oryzae* strain P131 expressing INV1-GFP fusion proteins, and this process was mediated by ALG3 (Fig. 4). Invertase is a major enzyme present in plants and microorganisms [29], and carries out the irreversible conversion of sucrose to glucose and fructose (**Fig. 6**A). A previous study showed that *M. oryzae* Δ*inv1* mutants had reduced fitness during plant infection [30]. INV1 has a length of 660 amino acids and contains 13 predicted *N*-glycosylation sites (Supplemental Table 2 and Fig. 5B). INV1 belongs to the glycosyl hydrolase family 32, containing a GH32 N-terminal domain, a GH32 C-terminal domain, and a signal peptide as a leading sequence (Fig. 6B).

**Figure 6.**
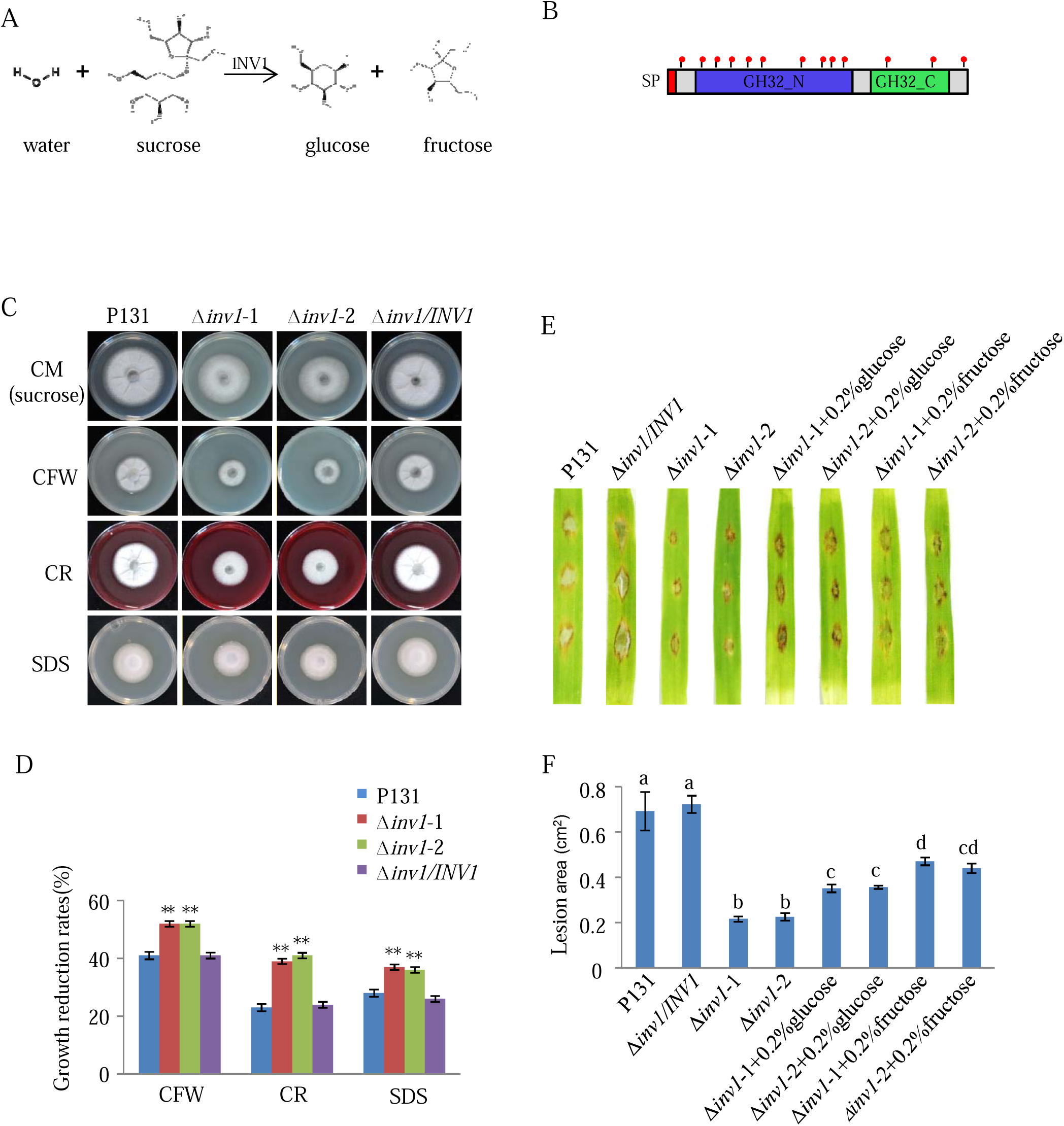
*INV1* is important for fungal virulence and cell wall integrity. **A**. The schematic diagram shows that INV1 functions in decomposition of sucrose. **B**. Deduced protein domains of INV1. Red dots indicate predicted *N*-glycosylation sites. **C**. Colony growth of strains P131, Δ*inv1-1*, Δ*inv1-2*, and Δ*inv1/INV1* on CM medium supplemented with the cell wall-disturbing agents 0.1 mg/ml CFW, 0.2 mg/ml Congo Red and 0.005%SDS. The cultures were incubated at 28[for 5 days before being photographed. **D**. Calculation of the growth reduction rates of mycelia growth on CM supplemented with cell wall-disturbing agents. Growth reduction rate (%) = [diameter (CM) − diameter (stress)]/diameter (CM). Error bars denote standard deviations from three biological replicates. **, statistically significant differences between the wild type and mutants (P<0.01, Student’s t test). **E**. Barley leaves drop-inoculated with conidium suspensions (2×10^4^ spores/mL) of the strains P131, Δ*inv1* and Δ*inv1/INV1* without or with exogenous glucose or fructose. The leaves were photographed at 5 dpi and the absolute lesion area was calculated (**F)**. Error bars denote standard deviations from three biological replicates with at least 9 leaves. The letters indicate significantly different statistical groups (P<0.05, one-way ANOVA with post-hoc Turkey tests) for corresponding strains or treatments.

We further characterized the function of INV1 in cell wall integrity and virulence by generating two independent knockout mutants in the P131 background, Δ*inv1-1* and Δ*inv1-2*, both mutants were confirmed with Southern blot analysis (Supplemental Fig. 3A and B), and we generated one complementation strain. The Δ*inv1* mutant strains grew normally on oatmeal-tomato agar plates (Supplemental Fig. 3C). To confirm its invertase function, the Δ*inv1* mutant strains were grown on solid MM without a carbon source or were supplied with sucrose, glucose, or fructose as the only carbon source. Since the size of the colonies was similar among the P131 strain and Δ*inv1* mutant strains, the thickness of the fungal growth reflects the carbon absorption efficiency. The mycelium of the P131 strain was very thin on the solid MM without a carbon source, but was much thicker on the solid MM with sucrose, glucose, or fructose. By contrast, the mycelium of the Δ*inv1* mutant strain was thicker only on the solid MM with glucose or fructose, but not with sucrose, confirming that the Δ*inv1* mutants could not utilize sucrose for fungal mycelium growth (Supplemental Fig. 3D). Introducing a functional copy of *INV1* into the Δ*inv1* mutant strain restored the sucrose utilization in the resulting Δ*inv1/INV1* complementation strain (Supplemental Fig. 3D). The fungal mycelial weights of Δ*inv1* mutant strains were also measured in the liquid MM supplied with different carbon sources and the same results were observed (Supplemental Fig. 3D). Therefore, we concluded that INV1 plays an important role in sucrose utilization.

In our previous report, we observed defects of cell wall integrity in the Δ*alg3* strains, but not in the Δ*slp1* mutant, suggesting that other secreted proteins regulated by ALG3 should function in cell wall integrity. Since INV1 is the most abundant protein secreted in CM during mycelium growth, we wondered whether INV1 plays a role in cell wall integrity. We grew Δ*inv1* mutant strains on common perturbing reagents used in cell wall integrity test, including Calcofluor white (CFW), Congo red (CR), and SDS [26]. When the P131 strain and the complementation strain Δ*inv1/INV1* were grown on CM with those agents, the vegetative growth was reduced by 41% with CFW, 23% with CR, and 28% with SDS, in comparison with their growth on CM. By contrast, growth in the mutant strains Δ*inv1-1* and Δ*inv1-2* was reduced by 52% with CFW, 39% with CR, and 37% with SDS, suggesting an increased sensitivity to cell wall perturbing reagents (Fig. 6C and D). We noticed that the sugar source commonly used in CM is sucrose, which cannot be utilized by the Δ*inv1* mutants. When we replaced the sucrose with glucose in this experiment, we found that all strains had similar reductions in growth rates on plates with CFW, CR, and SDS (Supplemental Fig. 3F and G). The growth assays in different CM revealed that fungal cell wall integrity is associated with normal carbon utilization in Δ*inv1* mutants.

We questioned whether the reduced pathogenicity of Δ*inv1* mutants was also associated with carbon uptake. Therefore, we investigated the role of INV1 on *M. oryzae* pathogenicity. Consistent with our previous report, the Δ*inv1* mutants showed significantly reduced virulence on rice and barley (*Hordeum vulgare*) leaves (Supplemental Fig. 3H, I, J, and K). Furthermore, we added an additional carbon source to see whether it could restore the mutant’s virulence. In drop infection assays, the symptom lesions from Δ*inv1* strains were much narrower than from the P131 strains on the unwounded barley leaves (Fig. 6E). When 0.2% glucose and 0.2% fructose were applied in the drop infection assays, the pathogenicity of Δ*inv1* mutants was partially restored (Fig. 6E and F). Therefore, carbon utilization is partially linked to the fungal virulence in Δ*inv1* mutants.

### MGG_04732 is important for fungal pathogenicity and cell wall integrity

We next tested the gene *MGG_04732*, encoding a protein with homologue to <u>A</u>cid <u>m</u>ammalian <u>c</u>hitin<u>ase</u> (named **AMCase** accordingly) containing a signal peptide followed with a glycosyl hydrolase family 18 (GH_18) domain (**Fig. 7**A) [31, 32]. Magnaporthe oryzae genome have 15 genes encoding GH_18 family chitinase, in which only *Chia1* was characterized to be important for fungal pathogenicity [33]. The *N*-glycosylation of AMCase was also confirmed in the mycelium proteins of P131 expressing AMCase -GFP fusion proteins, and this process was mediated by ALG3 (Fig. 4E).

**Figure 7.**
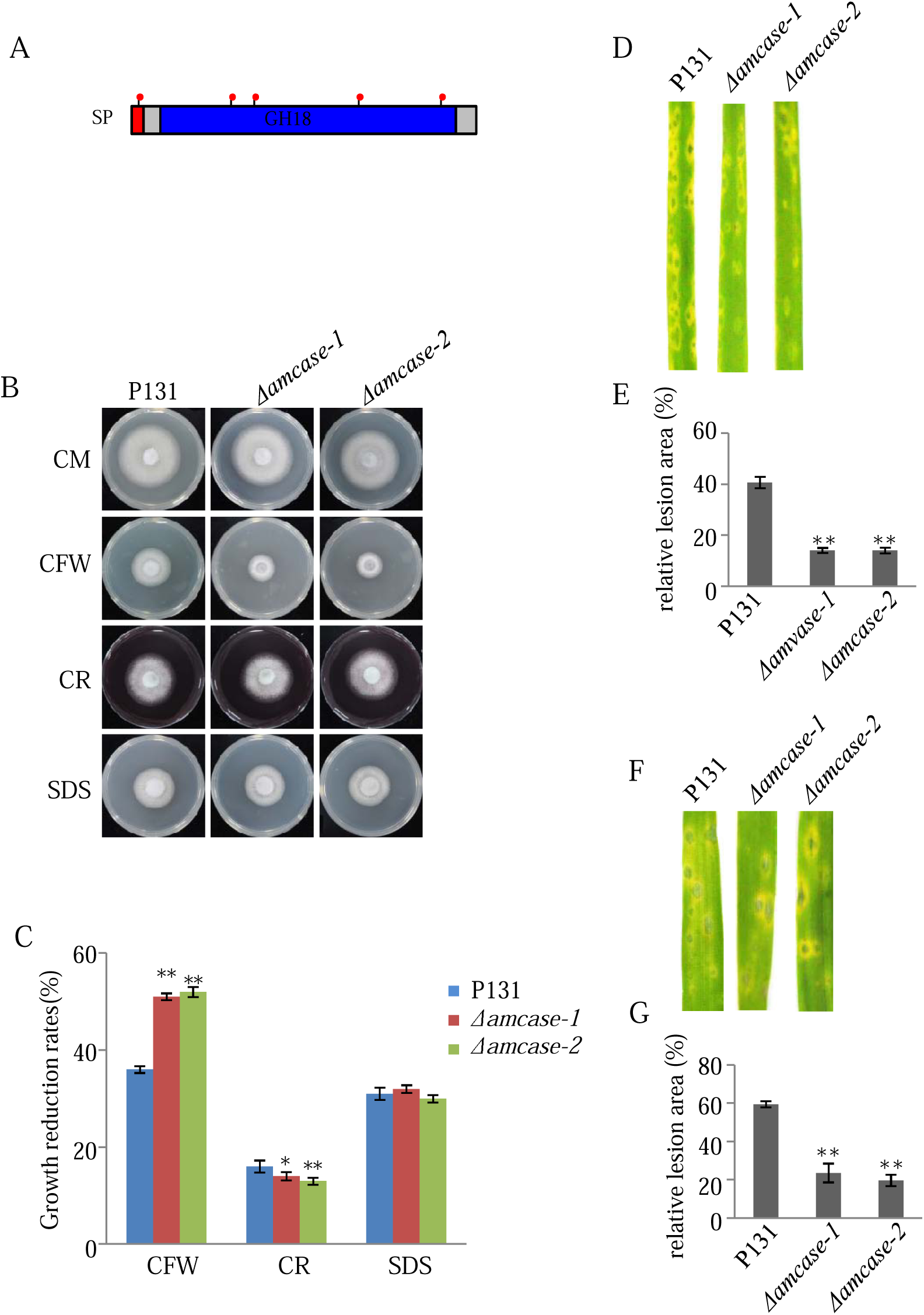
*AMCase* is important for fungal virulence and cell wall integrity. **A**. Deduced protein domains of AMCase. Red dots indicate predicted *N*-glycosylation sites. **B**. Colony growth of strains P131, *Δamcase-1, Δamcase-2*, and *Δamcase-3* on CM medium supplemented with the cell wall-disturbing agents 0.1 mg/ml CFW, 0.2 mg/ml Congo Red and 0.005%SDS. The cultures were incubated at 28°C for 5 days before being photographed. **C**. Calculation of the growth reduction rates of mycelia growth on CM supplemented with cell wall-disturbing agents. Growth reduction rate (%) = [diameter (CM) − diameter (stress)]/diameter (CM). Error bars denote standard deviations from three biological replicates. *, statistically significant differences between the wild type and mutants (P<0.05, Student’s t test). **, statistically significant differences between the wild type and mutants (P<0.01, Student’s t test). **D**. Rice leaves were sprayed with conidium suspensions (1×10^5^ spores/mL) of the strains P131, *Δamcase-1, Δamcase-2*, and *Δamcase-3*. The leaves were photographed at 5dpi and the relative lesion area was calculated (**E)**. Error bars denote standard deviations from three biological replicates with at least 9 leaves. **, statistically significant differences between the wild type and mutants (P<0.01, Student’s t test). **F**. Barley leaves were sprayed with conidium suspensions (1×10^4^ spores/mL) of the strains P131, *Δamcase-1, Δamcase-2*, and *Δamcase-3*. The leaves were photographed at 5dpi and the relative lesion area was calculated (**G)**. Error bars denote standard deviations from three biological replicates with at least 9 leaves. **, statistically significant differences between the wild type and mutants (P<0.01, Student’s t test).

We wondered whether *AMCase* also plays a role in cell wall integrity and pathogenicity, three independent knockout mutant *Δamcase-1, Δamcase-2* and Δ *amcase-3* were generated and confirmed by PCR analysis (Supplemental Fig. 4A and B). Those knockout mutant strains grew normal on the oatmeal-tomato agar (OTA) plates (Supplemental Fig. 4C). In the cell wall integration test, when the P131 strain was grown on CM with cell wall perturbing reagents, vegetative growth was reduced by 36% with CFW, 16% with CR, and 31% with SDS, in comparison with their growth on CM. By contrast, growth in the mutant strains *Δamcase-1, Δamcase-2*, and *Δamcase-3* was reduced by 50, 13, and 30% on plates with CFW, CR, and SDS, respectively, suggesting an increased sensitivity to CFW, but reduced sensitivity to CR (Fig. 7B and C). In the pathogenicity assay, we sprayed rice and barley leaves with spore solution. The knockout mutants showed significantly reduced pathogenecity on rice and barley (*Hordeum vulgare*) leaves (Fig. 7D, E, F and G), suggesting that AMCase functions in fungal pathogenicity. Since AMCase is related to the decomposition of chitin, we speculated AMCase might be also involved in chitin decomposition. We grew the wild type P131 and knockout strains on the solid minimal medium (MM) without carbon source or supplied with glucose and chitin as sole carbon source. The colony diameter of knockout strains slightly decreased compared with P131 on MM without carbon source, but no growth difference on MM supplied with glucose and chitin (Supplemental Fig 4F and G).

## Discussion

Many studies have used liquid culture medium to identify the secretome from pathogenic microbes, eg. *Xanthomonas oryzae, Botrytis cinerea*, and *Staphylococcus aureus* [34-37]. In the present study, we used a quantitative proteomics approach to analyze the secretome of *M. oryzae* and identified 558 proteins from the liquid culture medium, including 411 proteins of the wild-type P131 strain and 539 proteins of the Δ*alg3* mutant strain. Among the 558 proteins, 190 were found in previous *M. oryzae* secretome studies (Supplemental Table 2). In a previous study, 51 proteins were identified by comparative secretome analyses of *M. oryzae* grown in CM, MM, and MM-N (MM lacking N) media, of which 34 proteins (66.7%) were also identified in our data set [38]. In another study, the secretome of conidiospores grown in glass plates, PVDF membrane, and liquid culture medium were analyzed and 52 proteins were identified [39], of which 32 proteins (61.5%) overlapped with our secretome (Supplemental Table 2). Therefore, our *in vitro* secretome analysis detected many more secreted proteins compared with the two previous studies, possibly because we used gel-free rather than gel-based proteomics. Although it is a much greater challenge, *in vivo* apoplastic fluid of rice after *M. oryzae* infection was analyzed and a total of 441 secreted proteins were identified, of which 52 proteins were from a 2-DE gel-based approach and 425 proteins were from a gel-free MudPIT analysis [40]. When comparing this *in vivo* secretome with our *in vitro* secretome data, 171 proteins overlapped, accounting for 38.8–41.6% of the proteins identified in the two studies (Supplemental Table 2), indicating that a portion of pathogen infection-related proteins were expressed and secreted when grown in the liquid culture. Interestingly, many previously characterized secreted proteins were identified from our secretome dataset common in the P131 strain and the Δ*alg3* strain. Secreted proteins including MSP1 (MGG_05344), EMP1 (MGG_00527), MoHrip1 (MGG_15022), and MoHrip2 (MGG_16187) were found to be important for fungal pathogenicity [41-43]. Two secreted chitin deacylates MoCDA1 (MGG_14966), and MoCDA 2 (MGG_08774) were found to be dispensable for *Magnaporthe* virulence but important for fungal vegetative growth under stress conditions [44]. Notably, apart from EMP1, which contains four potential N-glycosylation sites, the other above-mentioned secreted proteins have only one or even no potential N-glycosylation sites, which could be explained as that N-glycosylation is dispensable for secretion of those proteins, thus they were detected both from P131 strain and the Δ*alg3* strain. Therefore, in the common list of proteins secreted from both strains, some uncharacterized secreted proteins might also function in fungal pathology or vegetative growth in different stress conditions. Further investigation of those commonly secreted proteins might identify additional virulence effectors in the future.

Protein *N*-glycosylation is a significant protein modification essential for fungal infection. Mutants of protein *N*-glycosylation showing defected in fungal virulence are reported not only for *M. oryzae* [45], but also for other plant pathogens such as *Penicilium digitatum, Botrytis cinerea* and *Ustilago maydis* [24, 25, 46-52]. Recent study on glycoprotein proteomics in *Ustilago maydis* using wildtype and mutants Δ*gls1*, a mutant also involved in *N*-glycosylation process, had identified several protein with *N*-glycosylation modification [53]. In our study, comparing the secretome of P131 and Δ*alg3*, 17 proteins were totally absent in the Δ*alg3* strain and 34 proteins had a significantly higher abundance in P131 samples than in Δ*alg3* samples; thus, these 51 proteins were considered secreted proteins that specifically require ALG3 for their secretion (Table 3). Since ALG3 functions in the early step of oligosaccharide synthesis for *N*-glycosylated proteins, the modified proteins should contain potential *N*-glycosylation sites. Among those 51 proteins, the 45 proteins that contain at least one predicted *N*-glycosylation site should be direct targets of ALG3 (Table 3); the other 6 proteins that do not have a predicted *N*-glycosylation site could be indirect targets of ALG3, as their secretion may be influenced by other ALG3-regulated proteins. It is worth to notice that our proteomics approach identified previously reported proteins involved in cell wall integrity, such as Gel3 (MGG_ 08370) and Gel4 (MGG_11861) [44]. Thus, a reduced secretion level of Gel3 and Gel4 in the Δ*alg3* mutant might account for the defects of cell wall integrity in Δ*alg3* mutants.

A subset of secreted proteins containing one predicted *N*-glycan site (protein encoded by MGG_09460), two *N*-glycan sites (MGG_13764), and more than three *N*-glycan sites (MGG_05785, MGG_01956, MGG_08772, MGG_03826, MGG_10209, and MGG_10466) were chosen for a series of validation assays. We confirmed that all candidate proteins indeed had *N*-glycosylation modifications (Fig. 4). The secretion test results suggest that all proteins were secreted in the growth medium. The secretion and localization of most of the tested proteins were affected in the Δ*alg3* mutant, validating that *N*-glycosylation is important for protein secretion and proper localization. Both the secretion and the localization of the protein encoded by MGG_13764 were not affected in the Δ*alg3* mutant, suggesting that for this specific protein, *N*-glycosylation is not necessary for its secretion. The validation tests on those selected proteins indicate that our *in vitro* comparative secretome analysis is a powerful approach to identify novel proteins that undergo *N*-glycosylation mediated by ALG3.

*M. oryzae* effectors consist of cytoplasmic effectors and apoplastic effectors [16]. Interestingly, apoplastic effectors Slp1 and Bas4 both contain predicted *N*-glycosylation sites, but none of the cytoplasmic effectors had a predicted *N*-glycosylation site. Thus, it is possible that *N*-glycosylation is a common feature of apoplastic effectors. We observed that all tested secreted proteins localized at the plant–fungus interface surrounding the infection hyphae (Fig. 5), which resembles the localization of Slp1, further supporting that the *N*-glycosylation modification is important for protein secretion via the conventional secretory pathway.

*M. oryzae* is a hemibiotrophic fungus, which, similar to other biotrophic fungi, depends on the nutrients provided by host plants at its early infection stage. Sugar is one of the most abundant nutrients a pathogen can get from living plants. Indeed, invertase homologs have been reported in several obligate biotrophic pathogens. The invertase from the rust fungus *Uromyces fabae* was expressed during infection and localized in the extrahaustorial matrix membrane and was important in breaking down sucrose into D-glucose and D-fructose [54]. In addition, a report on wheat stripe rust (*Puccinia striiformis f. sp. tritici*) revealed that the invertase gene *PsINV* plays a role in *Pst* pathogenicity [55]. The *M. oryzae* genome contains four genes encoding invertases, which were clustered in a subclade when compared with other fungal invertases, but only *INV1* was identified as an abundant protein in our secretome dataset. We found that *N*-glycosylation and the localization of INV1 were severely affected in the Δ*alg3* mutant; moreover, INV1 was also identified as secreted proteins in all the other *M. oryzae* secretome datasets. A previous report confirmed that a Δ*inv1* mutant showed impaired growth on sucrose-containing media, including impaired biomass formation and virulence [30]. In addition, we showed that the cell wall integrity was affected in Δ*inv1* mutants when grown in CM with sucrose, but this phenotype was completely restored when we replaced the sucrose with glucose. In a fungal virulence test, adding 0.2% glucose and 0.2% fructose also partially restored the pathogenicity of the Δ*inv1* mutants. Taken together, these observations show that carbon utilization is important for fungal cell wall integrity and virulence in *M. oryzae*.

Chitin, a homopolymer of 1, 4-β-linked N-acetyl-D-glucosamine (GlcNAc), is a structural component of fungal cell wall [56]. Chitin fragments are well-characterized elicitors that induce plant immune responses in many plant species [57, 58]. Chitinases are chitin-degrading enzymes that ubiquitous exit in a wide range of organisms including viruses, bacteria, fungi, insects, plant and animals [59]. *M. oryzae* has 15 genes that are annotated as GH_18 family chitinases. MoChia1 is involved in the fungal virulence through degrading chitin into small fragments, which would escape the recognition of plant immune receptors [33, 60]. Different chitinases had their specific expression in different cell types, *AMCase* maintains the highest expression level in appressorium among 15 chitinases, indicating *AMCase* is likely to be associated with fungal infection [33]. Indeed, our study exhibited that *Δamcase* mutants were less virulent on both rice and barley leaves, confirming that *AMCase* play an important role in pathogenicity. In addition, we showed that the cell wall integrity was affected in *Δamcase* mutants, but with different sensitivity in CFW and CR, indicating that the mechanism underline the cell wall remodeling in response to CFW differs to that to CR.

Notably, our secretome analysis did not identify the well characterized apoplastic effector protein Slp1, which is known to undergo *N*-glycosylation mediated by ALG3. Since our research was conducted on vegetative mycelium growth, the expression of Slp1 might be lower than that in the *in vivo* infection condition. Many pathogenicity-related genes, especially effectors, have no or a low level of expression during vegetative growth, and higher expression specifically during infection [61, 62]. Recent studies reported the functions of various transcription factors and epigenetic control in regulating effector expression in phytopathogenic fungi. For example, methylation of lysine 9 and/or lysine 27 of histone H3 (H3K9me3, H3K27me3) seems to be related with heterochromatin and effector gene silencing, and methylation of lysine 4 of histone H3 (H3K4me2) with euchromatin and effector gene expression [63-66]. Further proteomics studies using related histone modification mutants would enhance our understanding of secreted effector proteins in *M. oryzae*.

## Materials and Methods

### Fungal strains and Growth Conditions

*M. oryzae* wild-type strain P131 (field isolate) and the mutant strains Δ*alg3* which lacking the *α*-1, 3-mannosyltransferase were used in this study. The Δ*alg3* strain was generated and verified previously [26]. *M. oryzae* strains with different transformants generated in this study were listed in supplemental Table 3. All *M. oryzae* strains were maintained on oatmeal tomato agar (OTA) medium at 28°C. [67]. The liquid complete medium (CM) contains 6g/L yeast extract, 3g/L casein acid hydrolysate, 3g/L casein enzymatic hydrolysate and 10g/L sucrose. And the liquid minimal medium (MM) contains 6g/L NaNO_3_, 0.502g/L KCl, 0.502g/L MgSO_4_.7H_2_O, 1.52g/L KH_2_PO_4_, 1000 X trace element, 10g/L D-glucose, 1% thiamine and 0.05% biotin (PH 6.5). For cell wall integrity test, 5mm mycelial blocks of different strains were placed on CM agar with 0.1 mg/mL CFW (Sigma-Aldrich, Shanghai, China), 0.2 mg/mL CR (Sigma-Aldrich, Shanghai, China) and 0.005% SDS. For the carbohydrate supplement test, 5mm mycelial blocks of different strains were placed on solid minimal medium without or with different carbohydrate supplements at 28°C for 5 days. For the mycelial wet-weight measurement, the 0.1g mycelia were grown on liquid minimal medium without or with different carbohydrate supplements at 28°C for 1 day.

### Virulence Test

Rice (Oryza sativa cv LTH) seedlings at the third leaf stage and barley (cv E9) seedlings with 7-d-old were used for virulence test. Different M. oryzae strains with 10^5^/mL conidia suspension were used for spray inoculation as described previously [68]. Leaves were photographed at 5 days after inoculation and relative lesion area on leaves was calculated. To test the effect of exogenous glucose and fructose on infection of the Δ*inv1* mutant, 0.2% glucose and 0.2% fructose was added onto the fungal inoculation site at 18 hpi after the drop inoculation, and subsequently photographs and statistics of lesion area were taken at 5 days after inoculation.

### Secretome sample extraction

Secreted proteins were extracted from liquid medium. One gram of mycelia was collected and grown in liquid culture for 18 hours, then the liquid medium was filtrated through Miracloth (Merck millipore, Beijing, China) and centrifugated at 12,000rpm for 10 min. Two hundred-milliliters supernatants were collected and added with 12.5% (v/v) trichloroacetic acid at 4□ overnight to precipitate proteins. The pellets were collected after centrifuging for 30 min at 12,000rpm, washed twice with 100% acetone and dried. The protein samples were stored at −80 °C for further analyses.

### LC-MS/MS analyses and data processing

Proteins were digested using FASP method [69]. Peptides were separated with a Waters (Milford, MA, USA) nanoAcquity nano-HPLC. Mobile phases were 0.1% FA in water (A) and 0.1% FA in acetonitrile (B). We used linear gradient from 1% B to 35% B in 65 min. The trap column was Thermo Acclaim PepMap 100 (75 μm×20 mm, C18, 3 μm). The analytical column was homemade (100 μm×200 mm C18 stationary phase, Phenomenex, Aqua 3 μm 125Å). nanospray ESI-MS was performed on a Thermo Q Exactive high resolution mass spectrometer (Thermo Scientific, Waltham, MA, USA) in a data-dependent mode.

Raw MS files were subjected to MaxQuant software (Version 1.6.0.1) for protein identification and quantification as described before [70]. Peak list was generated by Andromeda which is a built-in engine in MaxQuant and searched against a *Magnaporthe oryzae* database (download from https://www.broadinstitute.org) augmented with the reversed sequence. Trypsin/P was set as the enzyme for digestion. Carbamidomethyl (C) was set as a fixed modification, while Oxidation (M) and Acetyl (protein N-term) were set as variable modifications. The maximum of missed cleavage was set as 2. Main search peptide tolerance and MS/MS match tolerance were both set as 20 ppm. False discovery rates of peptide and protein were both set at 1%.

### Vector Construction

To generate fluorescent protein fused with GFP, coding regions of candidate proteins were cloned into pGTN under a *MoEF* (MGG_03641) promoter [71]. The resulting constructs were digested with *EcoRI* and delivered into the protoplasts of P131 and Δ*alg3* strains as described [68]. Media with 400 g/ml neomycin (Ameresco, Ohio, USA) were used to select neomycin-resistant transformants.

To generate the gene replacement constructs, flanking sequences with 1.5-kb upstream and downstream of targeted genes (*INV1* and *AMCase*) were amplified using genomic DNA of P131. The two flanking sequences were cloned into pKOV21 [68] as the deletion vector pKO-*genes*. The resulting constructs were linearized by *NotI* and delivered into the protoplasts of P131 to generate deletion mutants, which were confirmed by southern blot hybridization.

### Fluorescence intensity and Subcellular localization Analysis

To analysis fluorescence intensity, the transformants of candidate proteins fused with GFP was grown in liquid CM cultured at 160 rpm for 22 h. Then fluorescence intensity of 100ul filtered medium and 0.1g mycelium proteins was measured with microplate reader (Molecular Devices i3x, Shanghai, China), respectively. To observe subcellular localization, these transformants of these validated proteins fused with GFP inoculated in barley leaves were observed with an epifluorescence microscope (Nikon, Toyko, Japan).

### Protein *N*-Glycosylation Analysis

For glycosylation analysis, total proteins were extracted from mycelia using the protein lysis buffer (20 mM Tris-HCl, pH 7.5, 150 mM NaCl, 0.5 mM EDTA, 0.5% NP-40). Proteins with or without PNGase F (NEB, USA) digestion were mixed with the loading buffer (250Mm Tris-HCl PH 6.8, 10% SDS, 0.5% bromophenol blue, 50% glycerol, 5%β-mercaptoethanol) for Western Blot analysis. Antibodies of anti-GFP (1:5000; ABclonal, Hubei, China) and anti-Actin (1:10,000; ABclonal, Hubei, China) were used to detect GFP fusion proteins and actin control, respectively.

## Authors’ contributions

XL, YP, JY and NL conceived the project; LQ, XL, NL, CY and RW undertook proteomic analysis; NL, XL, MH, DC, YZ, XW and GY conducted experiments; XL, YP, NL and LQ wrote the article.

## Acknowledgments

We thank Zhen Li for the technique support of LC-MS/MS analysis. This work was supported by the National Key Research and Development Plan [2016YFD0300703], Chinese Universities Scientific Fund [2018TC012], the Program for Changjiang Scholars and Innovative Research Team in University [IRT1042], and the 111 Project [B13006].

## Supplementary material

**Supplemental Figure 1 Secreted proteins from *Magnaporthe oryzae* isolate P131 and** Δ***alg3* mutant**

**A**. Western blot shows the secretion of Slp1-GFP from P131 in CM and MM liquid medium. Total secreted protein was used as a loading control. **B**. Silver staining of secreted protein samples from the CM liquid culture of P131 and Δ*alg3* strains. Three replicates were performed for each strain.

**Supplemental Figure 2.** The GFP signal from control samples.

**A**. Immunoblotting analysis of protein 03670-GFP, 10234-GFP and 10318-GFP (Group 3) from culture supernatants and mycelial extracts from wild-type P131 and Δ*alg3* strains. **B**. GFP fluorescent was measured in 96 well plates (upper panel) and the ratio of GFP intensity from CM liquid culture versus mycelial protein extracts was calculated (lower panel) for Slp1-GFP, MoGRP1-GFP, control GFP samples and background control P131 samples. Error bars denote standard deviations from three biological replicates. **, statistically significant differences between the wild type and mutants (P<0.01, Student’s t test). **C**. The fluorescence signal from control GFP samples. Left panel, Fluorescence signal is imaged in infected hyphae growing in barley epidermal cells for the tested proteins. White lines in the figures show the path for fluorescence intensity measurement. Middle panel, the brightfield image for the infection hyphae. Right panel, the fluorescence intensity measured using ImageJ. Scale bar=20

**Supplemental Figure 3 The characterization of *M. oryzae INV1* gene function**

**A**. Schematic diagram of the *INV1* deletion strategy. **B**. DNA gel blot analysis of the *INV1* deletion mutants. **C**. Colony growth of wild-type strain P131, *INV1* deletion mutant Δ*inv1-1* and Δ*inv1-2*, and complemented strain Δ*inv1/INV1* on OTA medium. **D**. *M. oryzae* stains P131, Δ*inv1-1*, Δ*inv1-2* and Δ*inv1*/INV were grown on minimal medium (MM) with different carbon sources (sucrose, glucose and fructose) and the picture were taken at day 5 grown in 28°C. **E**. The mycelia fresh weight of strains P131, Δ*inv1-1*, Δ*inv1-2* and Δ*inv1*/INV grown in MM liquid media was measured without or with different carbon sources for 24 hours. Error bars denote standard deviations from three biological replicates. **, statistically significant differences between the wild type and mutants (P<0.01, Student’s t test). **F**. Colony growth of strains P131, Δ*inv1-1*, Δ*inv1-2*, and Δ*inv1/INV1* on CM with glucose supplemented with the cell wall-disturbing agents 0.1 mg/ml CFW, 0.2 mg/ml Congo Red and 0.005%SDS. The cultures were incubated at 28[for 5 days before being photographed. **G**. Statistical analysis of the growth reduction rates of mycelia growth on CM with glucose supplemented with cell wall-disturbing agents. Growth reduction rate (%) = [diameter (CM) − diameter (stress)]/diameter (CM). Error bars denote standard deviations from three biological replicates. **H**. Rice leaves were sprayed with conidium suspensions (1×10^5^ spores/mL) of the strains P131, Δ*inv1-1*, Δ*inv1-2*, and Δ*inv1/INV1*. The leaves were photographed at 5dpi and the relative lesion area was calculated (**I)**. Error bars denote standard deviations from three biological replicates with at least 9 plants. **, statistically significant differences between the wild type and mutants (P<0.01, Student’s t test). **J**. Barley leaves were sprayed with conidium suspensions (1×10^4^ spores/mL) of the strains P131, Δ*inv1-1*, Δ*inv1-2*, and Δ*inv1/INV1*. The leaves were photographed at 5dpi and the relative lesion area was calculated (**K)**. Error bars denote standard deviations from three biological replicates with at least 9 plants. **, statistically significant differences between the wild type and mutants (P<0.01, Student’s t test).

**Supplemental Figure 4 The characterization of *M. oryzae AMCase* gene function**

**A**. Schematic diagram of the *AMCase* deletion strategy. **B**. DNA gel blot analysis of the *AMCase* deletion mutants. **C**. Colony growth of wild-type strain P131, *AMCase* deletion mutant *Δamcase-1, Δamcase-2* and *Δamcase -3* on OTA medium. **D**. The colony diameter was calculated for wild-type P131 and *AMCase* deletion mutants grown on OTA medium. **E**. Statistical analysis of the sporulation of wild-type P131 and *AMCase* deletion mutants. **F**. *M. oryzae* stains P131, *Δamcase-1, Δamcase-2* and *Δamcase-3* were grown on minimal medium (MM) with different carbon sources (glucose and chitin) and the picture were taken at day 5 grown in 28°C. **G**. The colony diameter was calculated for wild-type P131 and *AMCase* deletion mutants grown on minimal medium (MM) with different carbon sources. Error bars denote standard deviations from three biological replicates.

## Supplemental Tables

**Supplemental Table 1 The identified secretory proteins of P131and** Δ***alg3***

**Supplemental Table 2 Bioinformatics analysis identified secretory proteins**

**Supplemental Table 3 List of strains used in this study**

**Supplemental Table 4 List of oligonucleotide primers used in this study**

## References

[1] Dean R, Van Kan JA, Pretorius ZA, Hammond-Kosack KE, Di Pietro A, Spanu PD, et al. The Top 10 fungal pathogens in molecular plant pathology. Mol Plant Pathol 2012;13:414–30.

[2] Horbach R, Navarro-Quesada AR, Knogge W, Deising HB. When and how to kill a plant cell: Infection strategies of plant pathogenic fungi. Journal of Plant Physiology 2011;168:51–62.

[3] Kankanala P, Czymmek K, Valent B. Roles for rice membrane dynamics and plasmodesmata during biotrophic invasion by the blast fungus. Plant Cell 2007;19:706–24.

[4] Fernandez J, Orth K. Rise of a Cereal Killer: The Biology of Magnaporthe oryzae Biotrophic Growth. Trends Microbiol 2018;26:582–97.

[5] Dagdas YF, Yoshino K, Dagdas G, Ryder LS, Bielska E, Steinberg G, et al. Septin-mediated plant cell invasion by the rice blast fungus, Magnaporthe oryzae. Science 2012;336:1590–5.

[6] Egan MJ, Wang ZY, Jones MA, Smirnoff N, Talbot NJ. Generation of reactive oxygen species by fungal NADPH oxidases is required for rice blast disease. Proc Natl Acad Sci U S A 2007;104:11772–7.

[7] Foster AJ, Ryder LS, Kershaw MJ, Talbot NJ. The role of glycerol in the pathogenic lifestyle of the rice blast fungus Magnaporthe oryzae. Environ Microbiol 2017;19:1008–16.

[8] Mosquera G, Giraldo MC, Khang CH, Coughlan S, Valent B. Interaction Transcriptome Analysis Identifies Magnaporthe oryzae BAS1-4 as Biotrophy-Associated Secreted Proteins in Rice Blast Disease. Plant Cell 2009;21:1273–90.

[9] Ryder LS, Talbot NJ. Regulation of appressorium development in pathogenic fungi. Curr Opin Plant Biol 2015;26:8–13.

[10] Yi M, Valent B. Communication Between Filamentous Pathogens and Plants at the Biotrophic Interface. Annual Review of Phytopathology, Vol 51 2013;51:587–611.

[11] Chanclud E, Kisiala A, Emery NR, Chalvon V, Ducasse A, Romiti-Michel C, et al. Cytokinin Production by the Rice Blast Fungus Is a Pivotal Requirement for Full Virulence. PLoS Pathog 2016;12:e1005457.

[12] Mentlak TA, Kombrink A, Shinya T, Ryder LS, Otomo I, Saitoh H, et al. Effector-Mediated Suppression of Chitin-Triggered Immunity by Magnaporthe oryzae Is Necessary for Rice Blast Disease. Plant Cell 2012;24:322–35.

[13] Patkar RN, Benke PI, Qu Z, Chen YY, Yang F, Swarup S, et al. A fungal monooxygenase-derived jasmonate attenuates host innate immunity. Nat Chem Biol 2015;11:733–40.

[14] Skamnioti P, Gurr SJ. Magnaporthe grisea cutinase2 mediates appressorium differentiation and host penetration and is required for full virulence. Plant Cell 2007;19:2674–89.

[15] Van Vu B, Itoh K, Nguyen QB, Tosa Y, Nakayashiki H. Cellulases belonging to glycoside hydrolase families 6 and 7 contribute to the virulence of Magnaporthe oryzae. Mol Plant Microbe Interact 2012;25:1135–41.

[16] Giraldo MC, Dagdas YF, Gupta YK, Mentlak TA, Yi M, Martinez-Rocha AL, et al. Two distinct secretion systems facilitate tissue invasion by the rice blast fungus Magnaporthe oryzae. Nature Communications 2013;4.

[17] Cesari S, Thilliez G, Ribot C, Chalvon V, Michel C, Jauneau A, et al. The Rice Resistance Protein Pair RGA4/RGA5 Recognizes the Magnaporthe oryzae Effectors AVR-Pia and AVR1-CO39 by Direct Binding. Plant Cell 2013;25:1463–81.

[18] Jia Y, McAdams SA, Bryan GT, Hershey HP, Valent B. Direct interaction of resistance gene and avirulence gene products confers rice blast resistance. Embo Journal 2000;19:4004–14.

[19] Park CH, Chen S, Shirsekar G, Zhou B, Khang CH, Songkumarn P, et al. The Magnaporthe oryzae effector AvrPiz-t targets the RING E3 ubiquitin ligase APIP6 to suppress pathogen-associated molecular pattern-triggered immunity in rice. Plant Cell 2012;24:4748–62.

[20] Park CH, Shirsekar G, Bellizzi M, Chen S, Songkumarn P, Xie X, et al. The E3 Ligase APIP10 Connects the Effector AvrPiz-t to the NLR Receptor Piz-t in Rice. PLoS Pathog 2016;12:e1005529.

[21] Shi X, Long Y, He F, Zhang C, Wang R, Zhang T, et al. The fungal pathogen Magnaporthe oryzae suppresses innate immunity by modulating a host potassium channel. PLoS Pathog 2018;14:e1006878.

[22] Wang R, Ning Y, Shi X, He F, Zhang C, Fan J, et al. Immunity to Rice Blast Disease by Suppression of Effector-Triggered Necrosis. Curr Biol 2016;26:2399–411.

[23] Yoshida K, Saitoh H, Fujisawa S, Kanzaki H, Matsumura H, Yoshida K, et al. Association Genetics Reveals Three Novel Avirulence Genes from the Rice Blast Fungal Pathogen Magnaporthe oryzae. Plant Cell 2009;21:1573–91.

[24] Fernandez-Alvarez A, Elias-Villalobos A, Jimenez-Martin A, Marin-Menguiano M, Ibeas JI. Endoplasmic Reticulum Glucosidases and Protein Quality Control Factors Cooperate to Establish Biotrophy in Ustilago maydis. Plant Cell 2013;25:4676–90.

[25] Schirawski J, Bohnert HU, Steinberg G, Snetselaar K, Adamikowa L, Kahmann R. Endoplasmic reticulum glucosidase II is required for pathogenicity of Ustilago maydis. Plant Cell 2005;17:3532–43.

[26] Chen XL, Shi T, Yang J, Shi W, Gao X, Chen D, et al. N-glycosylation of effector proteins by an alpha-1,3-mannosyltransferase is required for the rice blast fungus to evade host innate immunity. Plant Cell 2014;26:1360–76.

[27] Gao X, Yin C, Liu X, Peng J, Chen D, He D, et al. A glycine-rich protein MoGrp1 functions as a novel splicing factor to regulate fungal virulence and growth in Magnaporthe oryzae. Phytopathology Research 2019;1.

[28] Szigeti M, Bondar J, Gjerde D, Keresztessy Z, Szekrenyes A, Guttman A. Rapid N-glycan release from glycoproteins using immobilized PNGase F microcolumns. J Chromatogr B Analyt Technol Biomed Life Sci 2016;1032:139–43.

[29] Roitsch T, Balibrea ME, Hofmann M, Proels R, Sinha AK. Extracellular invertase: key metabolic enzyme and PR protein. Journal of Experimental Botany 2003;54:513–24.

[30] Lindsay RJ, Kershaw MJ, Pawlowska BJ, Talbot NJ, Gudelj I. Harbouring public good mutants within a pathogen population can increase both fitness and virulence. Elife 2016;5.

[31] Boot RG, Blommaart EF, Swart E, Ghauharali-van der Vlugt K, Bijl N, Moe C, et al. Identification of a novel acidic mammalian chitinase distinct from chitotriosidase. J Biol Chem 2001;276:6770–8.

[32] Olland AM, Strand J, Presman E, Czerwinski R, Joseph-McCarthy D, Krykbaev R, et al. Triad of polar residues implicated in pH specificity of acidic mammalian chitinase. Protein Sci 2009;18:569–78.

[33] Han YJ, Song LL, Peng CL, Liu X, Liu LH, Zhang YH, et al. A Magnaporthe Chitinase Interacts with a Rice Jacalin-Related Lectin to Promote Host Colonization. Plant Physiology 2019;179:1416–30.

[34] Busche T, Hillion M, Van Loi V, Berg D, Walther B, Semmler T, et al. Comparative Secretome Analyses of Human and Zoonotic Staphylococcus aureus Isolates CC8, CC22, and CC398. Mol Cell Proteomics 2018;17:2412–33.

[35] Gonzalez-Fernandez R, Valero-Galvan J, Gomez-Galvez FJ, Jorrin-Novo JV. Unraveling the in vitro secretome of the phytopathogen Botrytis cinerea to understand the interaction with its hosts. Front Plant Sci 2015;6:839.

[36] Wang Y, Gupta R, Song W, Huh HH, Lee SE, Wu J, et al. Label-free quantitative secretome analysis of Xanthomonas oryzae pv. oryzae highlights the involvement of a novel cysteine protease in its pathogenicity. J Proteomics 2017;169:202–14.

[37] Wang Y, Kim SG, Wu J, Huh HH, Lee SJ, Rakwal R, et al. Secretome analysis of the rice bacterium Xanthomonas oryzae (Xoo) using in vitro and in planta systems. Proteomics 2013;13:1901–12.

[38] Wang Y, Wu J, Park ZY, Kim SG, Rakwal R, Agrawal GK, et al. Comparative Secretome Investigation of Magnaporthe oryzae Proteins Responsive to Nitrogen Starvation. Journal of Proteome Research 2011;10:3136–48.

[39] Jung YH, Jeong SH, Kim SH, Singh R, Lee JE, Cho YS, et al. Secretome analysis of Magnaporthe oryzae using in vitro systems. Proteomics 2012;12:878–900.

[40] Kim SG, Wang Y, Lee KH, Park ZY, Park J, Wu J, et al. In-depth insight into in vivo apoplastic secretome of rice-Magnaporthe oryzae interaction. Journal of Proteomics 2013;78:58–71.

[41] Ahn N, Kim S, Choi W, Im KH, Lee YH. Extracellular matrix protein gene, EMP1, is required for appressorium formation and pathogenicity of the rice blast fungus, Magnaporthe grisea. Molecules and Cells 2004;17:166–73.

[42] Nie HZ, Zhang L, Zhuang HQ, Shi WJ, Yang XF, Qiu DW, et al. The Secreted Protein MoHrip1 Is Necessary for the Virulence of Magnaporthe oryzae. International Journal of Molecular Sciences 2019;20.

[43] Nie HZ, Zhang L, Zhuang HQ, Yang XF, Qiu DW, Zeng H. Secreted protein MoHrip2 is required for full virulence of Magnaporthe oryzae and modulation of rice immunity. Applied Microbiology and Biotechnology 2019;103:6153–67.

[44] Sun DD, Cao HJ, Shi YK, Huang PY, Dong B, Liu XH, et al. The regulatory factor X protein MoRfx1 is required for development and pathogenicity in the rice blast fungus Magnaporthe oryzae. Molecular Plant Pathology 2017;18:1075–88.

[45] Chen XL, Liu C, Tang B, Ren Z, Wang GL, Liu W. Quantitative proteomics analysis reveals important roles of N-glycosylation on ER quality control system for development and pathogenesis in Magnaporthe oryzae. PLoS Pathog 2020;16:e1008355.

[46] Fernandez-Alvarez A, Elias-Villalobos A, Ibeas JI. The O-Mannosyltransferase PMT4 Is Essential for Normal Appressorium Formation and Penetration in Ustilago maydis. Plant Cell 2009;21:3397–412.

[47] Fernandez-Alvarez A, Marin-Menguiano M, Lanver D, Jimenez-Martin A, Elias-Villalobos A, Perez-Pulido AJ, et al. Identification of O-mannosylated Virulence Factors in Ustilago maydis. Plos Pathogens 2012;8.

[48] Gonzalez M, Brito N, Frias M, Gonzalez C. Botrytis cinerea Protein O-Mannosyltransferases Play Critical Roles in Morphogenesis, Growth, and Virulence. Plos One 2013;8.

[49] Guo M, Tan LY, Nie X, Zhu XL, Pan YM, Gao ZM. The Pmt2p-Mediated Protein O-Mannosylation Is Required for Morphogenesis, Adhesive Properties, Cell Wall Integrity and Full Virulence of Magnaporthe oryzae. Frontiers in Microbiology 2016;7.

[50] Harries E, Gandia M, Carmona L, Marcos JF. The Penicillium digitatum protein O-mannosyltransferase Pmt2 is required for cell wall integrity, conidiogenesis, virulence and sensitivity to the antifungal peptide PAF26. Molecular Plant Pathology 2015;16:748–61.

[51] Li MY, Liu XY, Liu ZX, Sun Y, Liu MX, Wang XL, et al. Glycoside Hydrolase MoGls2 Controls Asexual/Sexual Development, Cell Wall Integrity and Infectious Growth in the Rice Blast Fungus (vol 11, e0162243, 2016). Plos One 2017;12.

[52] Pan YM, Pan R, Tan LY, Zhang ZG, Guo M. Pleiotropic roles of O-mannosyltransferase MoPmt4 in development and pathogenicity of Magnaporthe oryzae (vol 65, pg 223, 2019). Current Genetics 2019;65:241-.

[53] Marin-Menguiano M, Moreno-Sanchez I, Barrales RR, Fernandez-Alvarez A, Ibeas JI. N-glycosylation of the protein disulfide isomerase Pdi1 ensures full Ustilago maydis virulence. PLoS Pathog 2019;15:e1007687.

[54] Voegele RT, Wirsel S, Moll U, Lechner M, Mendgen K. Cloning and characterization of a novel invertase from the obligate biotroph Uromyces fabae and analysis of expression patterns of host and pathogen invertases in the course of infection. Mol Plant Microbe Interact 2006;19:625–34.

[55] Chang Q, Liu J, Lin X, Hu S, Yang Y, Li D, et al. A unique invertase is important for sugar absorption of an obligate biotrophic pathogen during infection. New Phytol 2017;215:1548–61.

[56] Duo-Chuan L. Review of fungal chitinases. Mycopathologia 2006;161:345–60.

[57] Kaku H, Nishizawa Y, Ishii-Minami N, Akimoto-Tomiyama C, Dohmae N, Takio K, et al. Plant cells recognize chitin fragments for defense signaling through a plasma membrane receptor. Proceedings of the National Academy of Sciences of the United States of America 2006;103:11086–91.

[58] Shibuya N, Minami E. Oligosaccharide signalling for defence responses in plant. Physiological and Molecular Plant Pathology 2001;59:223–33.

[59] McGowan J, Fitzpatrick DA. Genomic, Network, and Phylogenetic Analysis of the Oomycete Effector Arsenal. mSphere 2017;2.

[60] Yang C, Yu Y, Huang J, Meng F, Pang J, Zhao Q, et al. Binding of the Magnaporthe oryzae Chitinase MoChia1 by a Rice Tetratricopeptide Repeat Protein Allows Free Chitin to Trigger Immune Responses. Plant Cell 2019;31:172–88.

[61] Hacquard S, Joly DL, Lin YC, Tisserant E, Feau N, Delaruelle C, et al. A comprehensive analysis of genes encoding small secreted proteins identifies candidate effectors in Melampsora larici-populina (poplar leaf rust). Mol Plant Microbe Interact 2012;25:279–93.

[62] Wang QQ, Han CZ, Ferreira AO, Yu XL, Ye WW, Tripathy S, et al. Transcriptional Programming and Functional Interactions within the Phytophthora sojae RXLR Effector Repertoire. Plant Cell 2011;23:2064–86.

[63] Chujo T, Scott B. Histone H3K9 and H3K27 methylation regulates fungal alkaloid biosynthesis in a fungal endophyte-plant symbiosis. Mol Microbiol 2014;92:413–34.

[64] Connolly LR, Smith KM, Freitag M. The Fusarium graminearum histone H3 K27 methyltransferase KMT6 regulates development and expression of secondary metabolite gene clusters. PLoS Genet 2013;9:e1003916.

[65] Soyer JL, El Ghalid M, Glaser N, Ollivier B, Linglin J, Grandaubert J, et al. Epigenetic control of effector gene expression in the plant pathogenic fungus Leptosphaeria maculans. PLoS Genet 2014;10:e1004227.

[66] Wiemann P, Sieber CM, von Bargen KW, Studt L, Niehaus EM, Espino JJ, et al. Deciphering the cryptic genome: genome-wide analyses of the rice pathogen Fusarium fujikuroi reveal complex regulation of secondary metabolism and novel metabolites. PLoS Pathog 2013;9:e1003475.

[67] Peng YL, Shishiyama J. Temporal Sequence of Cytological Events in Rice Leaves Infected with Pyricularia-Oryzae. Canadian Journal of Botany-Revue Canadienne De Botanique 1988;66:730–5.

[68] Yang J, Zhao XY, Sun J, Kang ZS, Ding SL, Xu JR, et al. A Novel Protein Com1 Is Required for Normal Conidium Morphology and Full Virulence in Magnaporthe oryzae. Molecular Plant-Microbe Interactions 2010;23:112–23.

[69] Wisniewski JR, Zougman A, Nagaraj N, Mann M. Universal sample preparation method for proteome analysis. Nature Methods 2009;6:359–U60.

[70] Cox J, Mann M. MaxQuant enables high peptide identification rates, individualized p.p.b.-range mass accuracies and proteome-wide protein quantification. Nat Biotechnol 2008;26:1367–72.

[71] Skamnioti P, Gurr SJ. Magnaporthe grisea cutinase2 mediates appressorium differentiation and host penetration and is required for full virulence. Plant Cell 2007;19:2674–89.

