## Supplemental figure 1-4 for "Comparative secretome of *Magnaporthe oryzae* identified proteins involved in virulence and cell wall integrity"

### Slide 1
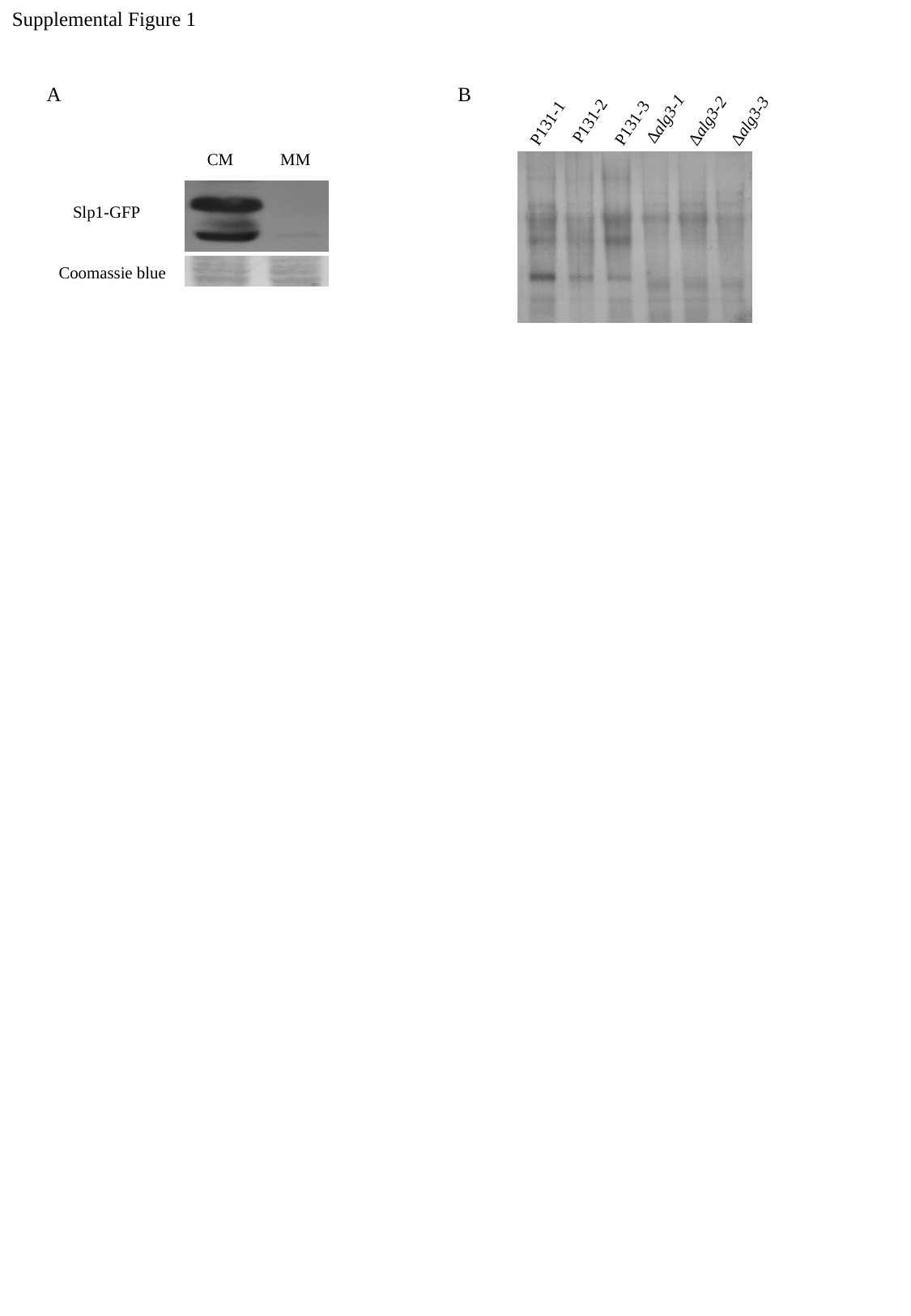

Supplemental Figure 1
A
B
∆alg3-1
∆alg3-2
∆alg3-3
P131-2
P131-3
P131-1
CM
MM
Slp1-GFP
Coomassie blue

### Slide 2
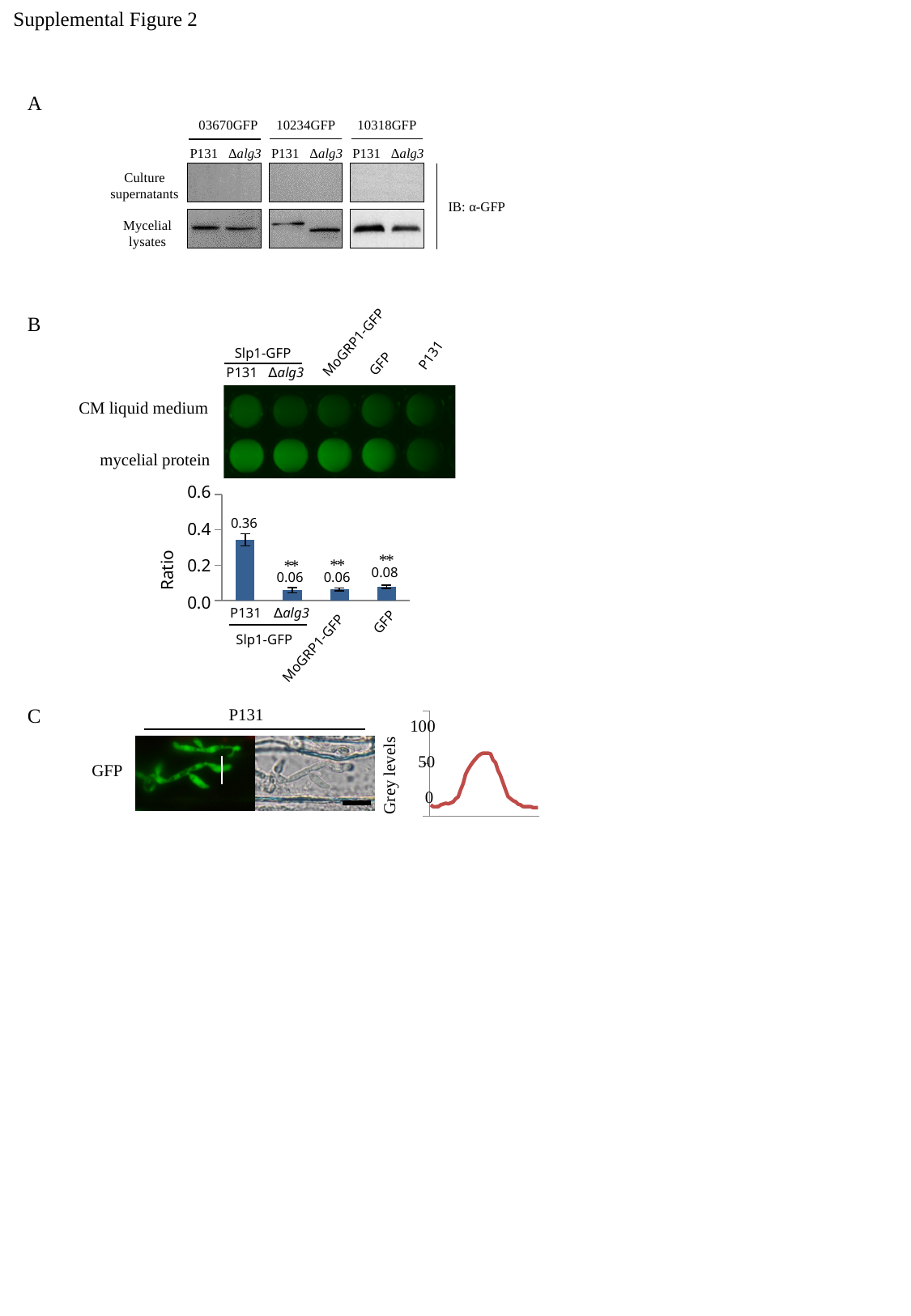

Supplemental Figure 2
A
10318GFP
P131
∆alg3
10234GFP
P131
∆alg3
03670GFP
P131
∆alg3
Culture supernatants
Mycelial
 lysates
IB: α-GFP
MoGRP1-GFP
Slp1-GFP
GFP
CM liquid medium
mycelial protein
0.6
#### Chart
| Category | |
|---|---|
| slp131 | 0.3438403893878028 |
| slpko | 0.058998203484748046 |
| 7511gfp | 0.063402066452388 |
| gfp | 0.0775157004451922 |0.4
0.2
0.0
P131
P131
∆alg3
0.36
Ratio
0.08
0.06
0.06
P131
∆alg3
GFP
Slp1-GFP
MoGRP1-GFP
B
 *
 *
 *
 *
 *
 *
C
P131
#### Chart
| Category | |
|---|---|
| 0 | 8.6667 |
| 3.3E-3 | 7.3333 |
| 6.7000000000000002E-3 | 7.3333 |
| 0.01 | 7.3798 |
| 1.3299999999999999E-2 | 8.7287 |
| 1.67E-2 | 9.3333 |
| 0.02 | 10.0 |
| 2.3300000000000001E-2 | 9.5504 |
| 2.6700000000000002E-2 | 10.124 |
| 0.03 | 11.0155 |
| 3.3300000000000003E-2 | 13.3566 |
| 3.6700000000000003E-2 | 14.7597 |
| 0.04 | 20.1318 |
| 4.3299999999999998E-2 | 24.8372 |
| 4.6699999999999998E-2 | 31.9922 |
| 0.05 | 35.6822 |
| 5.33E-2 | 38.7519 |
| 5.67E-2 | 41.4031 |
| 0.06 | 43.6667 |
| 6.3299999999999995E-2 | 45.7752 |
| 6.6699999999999995E-2 | 47.2248 |
| 7.0000000000000007E-2 | 48.0 |
| 7.3300000000000004E-2 | 48.0 |
| 7.6700000000000004E-2 | 48.0 |
| 0.08 | 47.4419 |
| 8.3299999999999999E-2 | 42.7829 |
| 8.6699999999999999E-2 | 40.7209 |
| 0.09 | 34.6589 |
| 9.3299999999999994E-2 | 30.7674 |
| 9.6699999999999994E-2 | 25.3256 |
| 0.1 | 20.1705 |
| 0.1033 | 14.9845 |
| 0.1067 | 13.4961 |
| 0.11 | 11.9457 |
| 0.1133 | 11.1938 |
| 0.1167 | 9.3333 |
| 0.12 | 8.7752 |
| 0.12330000000000001 | 7.4264 |
| 0.12670000000000001 | 7.4109 |
| 0.13 | 7.3333 |
| 0.1333 | 7.4264 |
| 0.13669999999999999 | 6.6667 |
| 0.14000000000000001 | 6.6512 |
| 0.14330000000000001 | 6.6667 |100
Grey levels
50
GFP
0

### Slide 3
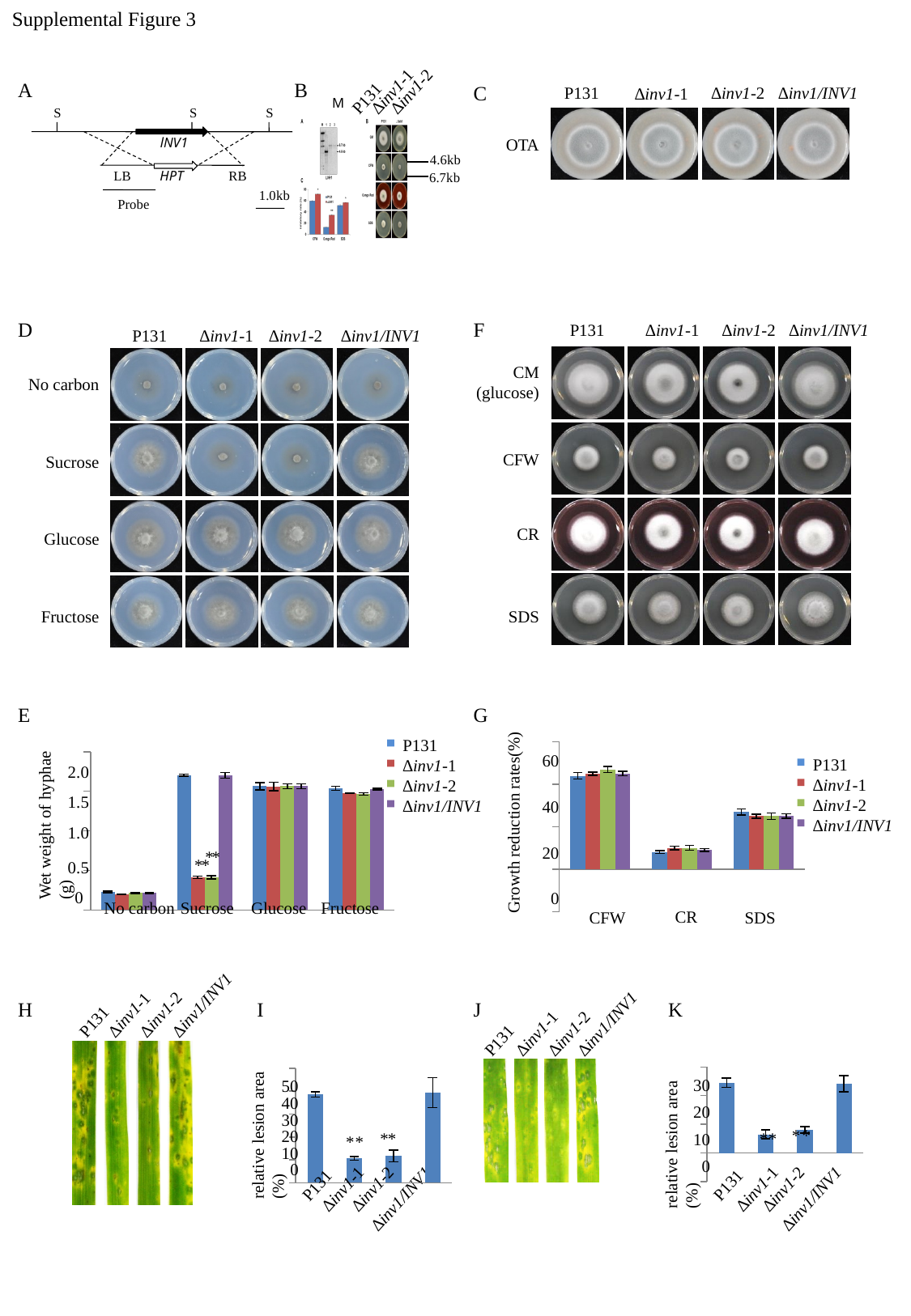

Supplemental Figure 3
∆inv1-1
∆inv1-2
P131
M
A
B
C
∆inv1-2
P131
∆inv1/INV1
∆inv1-1
S
S
S
LB
RB
1.0kb
Probe
lNV1
HPT
OTA
4.6kb
6.7kb
D
F
P131
∆inv1-1
∆inv1-2
∆inv1/INV1
P131
∆inv1-1
∆inv1-2
∆inv1/INV1
CM
(glucose)
No carbon
CFW
Sucrose
CR
Glucose
Fructose
SDS
E
G
Growth reduction rates(%)
#### Chart
| Category | P131 | 5785KO1 | 5785KO2 | 5785GFP |
|---|---|---|---|---|
| CFW | 0.44 | 0.45 | 0.47 | 0.45 |
| CR | 0.08 | 0.1 | 0.1 | 0.09 |
| SDS | 0.27 | 0.25 | 0.25 | 0.25 |60
P131
∆inv1-1
∆inv1-2
∆inv1/INV1
40
20
0
CR
SDS
CFW
P131
∆inv1-1
∆inv1-2
∆inv1/INV1
#### Chart
| Category | P131 | MoInvKO1 | MoInvKO2 | cMoInv |
|---|---|---|---|---|
| no carbon | 0.22999999999999998 | 0.20000000000000004 | 0.21333333333333335 | 0.21333333333333335 |
| sucrose | 1.7033333333333331 | 0.41333333333333333 | 0.41333333333333333 | 1.7 |
| glucose | 1.5633333333333332 | 1.5600000000000003 | 1.5633333333333335 | 1.5633333333333335 |
| fructose | 1.5366666666666664 | 1.4766666666666666 | 1.4666666666666666 | 1.5266666666666666 |2.0
1.5
1.0
0.5
0
No carbon
Sucrose
Glucose
Fructose
Wet weight of hyphae (g)
 *
 *
 *
 *
∆inv1/INV1
∆inv1-1
∆inv1-2
P131
∆inv1/INV1
∆inv1-1
∆inv1-2
P131
H
I
J
K
relative lesion area (%)
#### Chart
| Category | |
|---|---|
| P131 | 0.38556184195439785 |
| KO1 | 0.10646776800960855 |
| KO2 | 0.11702411834193077 |
| GFP | 0.3941927908052521 |50
40
30
20
10
0
P131
∆inv1-2
∆inv1-1
∆inv1/INV1
relative lesion area (%)
#### Chart
| Category | |
|---|---|
| P131 | 0.24507097190987268 |
| KO1 | 0.0649656567872793 |
| KO2 | 0.0804947616696835 |
| GFP | 0.2416046538557646 |30
20
10
0
P131
∆inv1-2
∆inv1-1
∆inv1/INV1
*
*
*
*
*
*
*
*

### Slide 4
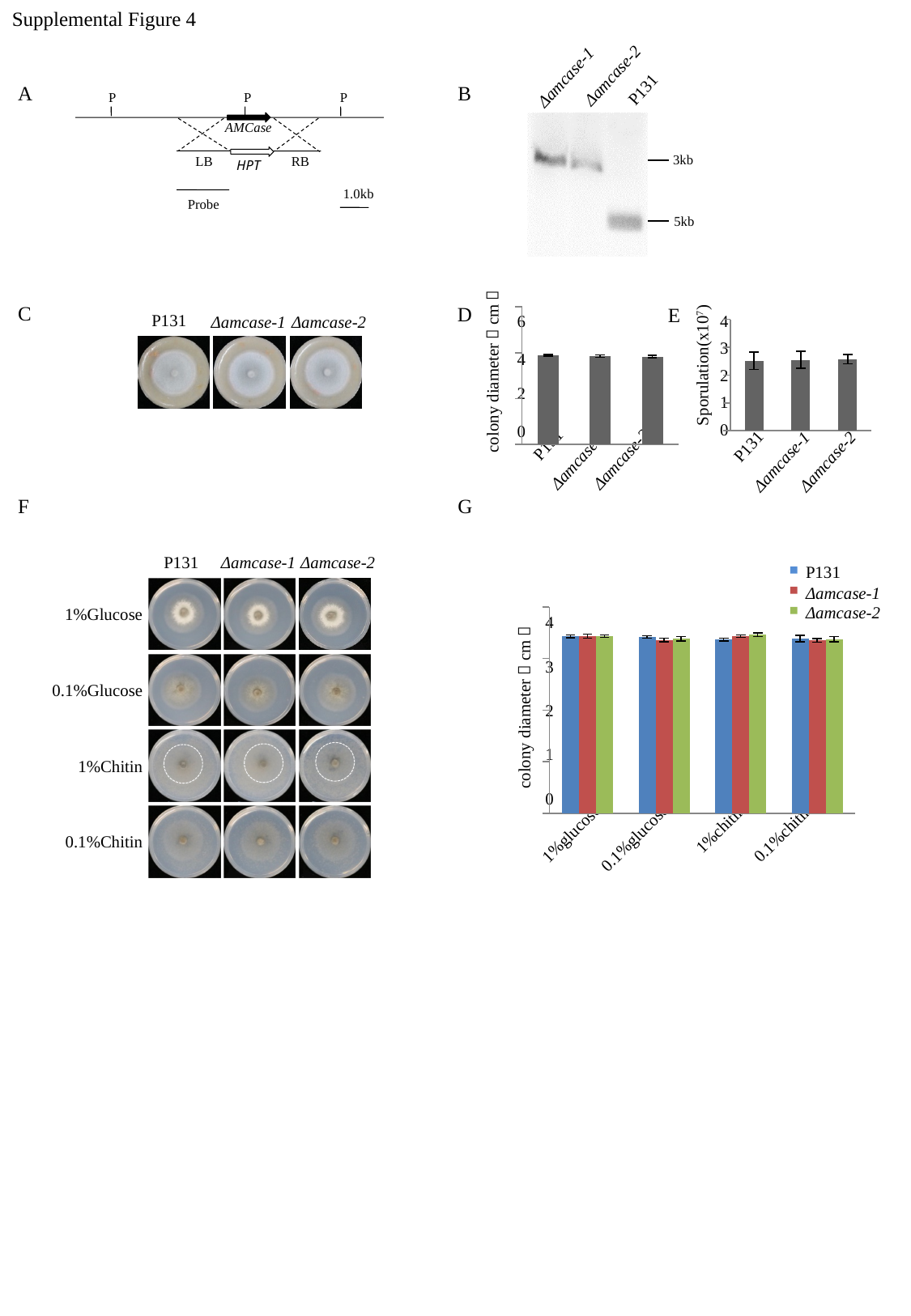

Supplemental Figure 4
Δamcase-2
Δamcase-1
P131
A
B
P
P
P
AMCase
3kb
LB
RB
HPT
1.0kb
Probe
5kb
C
D
E
P131
#### Chart
| Category | |
|---|---|
| P131 | 3.883333333333334 |
| KO1 | 3.85 |
| KO2 | 3.8249999999999997 |6
4
Δamcase-1
Δamcase-2
#### Chart
| Category | |
|---|---|
| P131 | 2.515 |
| KO1 | 2.5500000000000003 |
| KO2 | 2.5749999999999997 |3
4
Sporulation(x10^7^)
colony diameter（cm）
2
2
1
0
0
P131
P131
Δamcase-2
Δamcase-1
Δamcase-2
Δamcase-1
F
G
P131
Δamcase-1
Δamcase-2
P131
Δamcase-1
Δamcase-2
1%Glucose
#### Chart
| Category | P131 | 4732KO1 | 4732KO2 |
|---|---|---|---|
| 1% glucose | 3.4249999999999994 | 3.433333333333333 | 3.433333333333333 |
| 0.1% glucose | 3.416666666666666 | 3.358333333333334 | 3.383333333333333 |
| 1%chitin | 3.3666666666666667 | 3.433333333333333 | 3.4583333333333335 |
| 0.1%chitin | 3.3833333333333333 | 3.35 | 3.375 |4
3
0.1%Glucose
colony diameter（cm）
2
1
1%Chitin
0
1%chitin
0.1%chitin
1%glucose
0.1%glucose
0.1%Chitin
