## Supplemental table 3 for "Comparative secretome of *Magnaporthe oryzae* identified proteins involved in virulence and cell wall integrity"

**Supplemental Table 3 List of strains used in this study**

| **Strains** | **Genotype description** | **Reference** |
| --- | --- | --- |
| P131 | Wild type | Chen et al., 2014 |
| *Δalg3* | ALG3 deletion mutant of P131 | Chen et al., 2014 |
| MoGRP1GFP | MoGRP1-GFP transformant of P131 | Gao et al., 2019 |
| *ΔInv1* | Inv1 deletion mutant of P131 | This study |
| *ΔInv1/INV1* | Inv1 complement of *ΔInv1* | This study |
| *ΔAMCase1* | AMCase1 deletion mutant of P131 | This study |
| Slp1GFP | Slp-GFP transformant of P131 | This study |
| 05785GFP | 05785-GFP transformant of P131 | This study |
| 01956GFP | 01956-GFP transformant of P131 | This study |
| 08772GFP | 08772-GFP transformant of P131 | This study |
| 03826GFP | 03826-GFP transformant of P131 | This study |
| 09460GFP | 09460-GFP transformant of P131 | This study |
| 10209GFP | 10209-GFP transformant of P131 | This study |
| 10466GFP | 10466-GFP transformant of P131 | This study |
| 13764GFP | 13764-GFP transformant of P131 | This study |
| 00592GFP | 00592-GFP transformant of P131 | This study |
| 04732GFP | 04732-GFP transformant of P131 | This study |
| 03670GFP | 03670-GFP transformant of P132 | This study |
| 10234GFP | 10234-GFP transformant of P133 | This study |
| 10318GFP | 10318-GFP transformant of P134 | This study |
