## Supplemental table 4 for "Comparative secretome of *Magnaporthe oryzae* identified proteins involved in virulence and cell wall integrity"

**Supplemental Table 4** List of oligonucleotide primers used in this study

| **Name** | **Sequence (5’- 3’)** | **Applications** |
| --- | --- | --- |
| 03641promoter F | CGGAATTCTGGCTCTGGATACTGAAGGC | 03641promoter |
| 03641promoter R | CGGGATCCTTTGGCGGTTTGGTGCTC | 03641promoter |
| 05785F | caccaaaccgccaaaggatccATGAAATTCACATTTGTGTCATCGG | 05785-GFP |
| 05785R | acctctagaactagtggatccCCATCCGTTCCACCAGGTG | 05785-GFP |
| 01956F | caccaaaccgccaaaggatccATGAAGGCTACCTTTGTTACCCTC | 01956-GFP |
| 01956R | acctctagaactagtggatccCTGGAGTGTGACCTTTCCGC | 01956-GFP |
| 08772F | caccaaaccgccaaaggatccATGAAGGCAACAATCCTTACACTG | 08772-GFP |
| 08772R | acctctagaactagtggatccCAACAACATAGTAAGC | 08772-GFP |
| 03826F | caccaaaccgccaaaggatccATGCATCATCGCCGCCAT | 03826-GFP |
| 03826R | acctctagaactagtggatccAACCACTCTCAACACGGCATTAC | 03826-GFP |
| 07949F | caccaaaccgccaaaggatccATGGGCTACTACACCGACCACC | 07949-GFP |
| 07949R | acctctagaactagtggatccATCGCTGTCAGAGTCATTGTCG | 07949-GFP |
| 09460F | caccaaaccgccaaaggatccATGCAGCTCTCAAGCCTCCTC | 09460-GFP |
| 09460R | acctctagaactagtggatccAGTGGGTATGTAATGACAATGTCTGTT | 09460-GFP |
| 10209F | caccaaaccgccaaaggatccATGTTCTTCTTCAAAGTCTTGGTCG | 10209-GFP |
| 10209R | acctctagaactagtggatccCTTGACAAACTTGTACTGAACCGTC | 10209-GFP |
| 10466F | caccaaaccgccaaaggatccATGAAGGCTACGACGCTTTACAT | 10466-GFP |
| 10466R | acctctagaactagtggatccTCCGCTGCAACTGAATTCG | 10466-GFP |
| 13764F | caccaaaccgccaaaggatccATGCGTTCCTCGGTTGGTT | 13764-GFP |
| 13764R | acctctagaactagtggatccCTGACGCGGCATGTAGCG | 13764-GFP |
| 00592F | caccaaaccgccaaaggatccATGTACACAAAGACCAT | 00592-GFP |
| 00592R | acctctagaactagtggatccCAGCATGAAGTAACCCA | 00592-GFP |
| 04732F | caccaaaccgccaaaggatccATGTCACTCGTTAACCTCTCCAGG | 04732-GFP |
| 04732R | acctctagaactagtggatccCAGACCAACGGATCCGCG | 04732-GFP |
| 05785LBF | GGACTAGTGGTTTGAGTCATTGGCATC | Inv1 Knockout |
| 05785LBR | CGGAATTCGGGAGCGGAACAGATTACA | Inv1 Knockout |
| 05785RBF | CCATCGATCAGTCTCACTTGTGTCGTTGTC | Inv1 Knockout |
| 05785RBR | AAGGGCCCGCCAGAAGAAGTAAGGTGAAGT | Inv1 Knockout |
| 04732LBF | CGGGATCCCCCTTTCCAAGCCATTCCGT | AMCase Knockout |
| 04732LBR | CGGAATTCCGATTTGCACACTTGGGATGG | AMCase Knockout |
| 04732RBF | CCCAAGCTTTGGCGCAAGTTCGTCTGTAA | AMCase Knockout |
| 04732RBR | CCATCGATGCACCTCCTCGTTGTACCAG | AMCase Knockout |
| 04732UP | TAACCGCCCCTTTGTTAGCC | AMCase L check |
| HPT-UP | GACAGACGTCGCGGTGAGTT | AMCase L check |
| 04732DOWN | GTGCGTCCAAACACCTTGTA | AMCase R check |
| HPT-DOWN | TCTGGACCGATGGCTGTGTAG | AMCase R check |
| 04732 check F | ATGTCACTCGTTAACCTC | AMCase gene check |
| 04732 chevk R | CAGACCAACGGATCCGCG | AMCase gene check |
